# Oligodendroglia vulnerability in the human dorsal striatum in Parkinson’s disease

**DOI:** 10.1101/2025.01.15.633167

**Authors:** Juan M. Barba-Reyes, Lisbeth Harder, Sergio Marco Salas, Methasit Jaisa-aad, Clara Muñoz-Castro, Nima Rafati, Mats Nilsson, Bradley T. Hyman, Alberto Serrano-Pozo, Ana B. Muñoz-Manchado

**Affiliations:** Unit of Cell Biology, Department of Neuroscience. University of Cádiz. Institute for Biomedical Research and Innovation of Cádiz (INiBICA). Spain.; Laboratory of Molecular Neurobiology, Karolinska Institutet, Department of Medical Biochemistry and Biophysics, Stockholm, Sweden; Science for Life Laboratory, Department of Biochemistry and Biophysics, Stockholm University, Stockholm, Sweden; Institute of Computational Biology, Computational Health Center, Helmholtz, Munich; Department of Neurology, Massachusetts General Hospital, Boston, Massachusetts, USA; Harvard Medical School, Boston, Massachusetts, USA; National Bioinformatics Infrastructure Sweden, Uppsala University, Science for Life Laboratory, Department of Medical Biochemistry and Microbiology, Uppsala, Sweden

**Keywords:** Neurodegeneration, scRNA-seq, spatial transcriptomics, striatum, oligodendrocyte, myelin

## Abstract

Oligodendroglia are the responsible cells for myelination in the central nervous system and their involvement in Parkinson’s disease (PD) is poorly understood. We performed snRNA-seq and image-based spatial transcriptomics of human caudate nucleus and putamen (dorsal striatum) from PD and Control brain donors to elucidate the diversity of oligodendroglia and how they are affected by the disease. We have defined fifteen subclasses, from precursor to mature cells, four of which are disease-associated. These PD-specific populations are characterized by the overexpression of heat shock proteins and distinct expression signatures, including immune responses and myelination alterations. We have also identified disruptions in cell communication and oligodendrocyte development, evidenced by changes in neurotransmitter receptors expression and cell adhesion molecules. These transcriptomic changes correlated with impaired myelin integrity and altered oligodendrocyte distribution in the striatum. Thus, we uncover oligodendroglia as a critical cell type in PD and a potential new therapeutic target.

## Introduction

Parkinson’s disease (PD) has a significant impact on society due to its widespread implications both in patients’ health and caretakers, with a rapid increase in incidence over the past two decades^1,2^. PD is a heterogeneous condition, with mutations in multiple genes presenting with a similar parkinsonian syndrome and a single causative mutation resulting in a range of clinical presentations^3,4^. This heterogeneity suggests different underlying mechanisms at a biological level that are not yet fully understood. While the loss of dopaminergic neurons in the *substantia nigra pars compacta* (SNpc) and the accumulation of intracellular α-synuclein aggregates in Lewy bodies (LBs) and Lewy neurites (LNs) have long been associated with the development of PD^5,6^, recent studies have demonstrated the involvement of glial cells, in particular oligodendrocytes (OLs), in neurodegenerative disorders such as PD and Alzheimer’s disease (AD)^7–15^.

While they have traditionally been considered homogeneous populations^16–18^, recent advancements including single-cell/single-nucleus RNA-sequencing (sc/sn-RNA-seq) have revealed that OLs are heterogeneous, comprising numerous subpopulations and states that participate in distinct biological processes^19,20^, redefining the maturation process as a continuum. Indeed, OLs are known to progress through distinct developmental states, starting as oligodendrocyte progenitor cells (OPCs), which proliferate and differentiate into pre-myelinating OLs, that eventually mature into fully myelinating oligodendrocytes^21,22^. These different states correspond to distinct transcriptomic and morphologic profiles that can also vary along brain regions^23–28^.

Recent works suggest that alterations in myelin integrity and OL survival can play a crucial role in the progression of neurodegenerative diseases and multiple sclerosis (MS)^13–15,19,20,29^. Interestingly a few studies have also demonstrated a connection between OLs and PD^30–33^. Bryois *et al*^32^ found a genetic association between PD GWAS variants and OLs, while Wakabayashi *et al*^30^ observed α-synuclein inclusions in OLs of PD brains. These findings suggest that OLs may play a critical role in the development of PD. However, few studies have investigated the role of the OL lineage in PD, particularly in the dorsal striatum^31,34^, a brain region that includes the putamen (Pu) and caudate nucleus (CN) and is central to PD pathophysiology^35^ and therapeutic development, as it receives the nigrostriatal axons from the dopaminergic neurons of the SNpc^36^.

In this study, we elucidated the transcriptomic changes induced by PD across OLs lineage populations using snRNA-seq in the dorsal striatum from 63 donors and image-based spatial transcriptomics in a subset of them. Our results reveal distinct transcriptomic states of PD-associated oligodendrocytes (PDA-OLs), delineating diverse altered functional pathways, the spatial organization of OLs, and the potential biological consequences of OL dysfunction for the extent of both dopaminergic neuron loss and LB/LN formation.

## Materials and methods

### Human tissue

Human brain samples for this study were obtained from donors with a clinical and neuropathological diagnosis of PD and controls not meeting clinical or neuropathological diagnostic criteria for any neurodegenerative disease. Specifically, the CN and Pu were dissected from coronal slabs at the level of the *nucleus accumbens*. The Control group comprised 27 individuals (Pu = 27, CN = 24) ranging in age from 25 to over 90 years. The PD group included 36 individuals (Pu = 35, CN = 34) aged between 60 and over 90 years. Samples were collected from three sources: the Human Brain and Spinal Fluid Resource Center (Los Angeles, CA, USA), the Parkinson’s UK Brain Bank at Imperial College London (London, UK), and the Massachusetts Alzheimer’s Disease Research Center (Charlestown, MA, USA). Donors or their next-of-kin provided written informed consent for brain autopsy, and the study was approved by the review board of each brain bank. Detailed information regarding the samples is provided in Supplementary Table 1. Since PD is more common in men, the Control and PD groups were not matched by sex (i.e., there were 19 females and 44 males in the full cohort, Control: X/Y and PD: W/Z male/female). Consequently, the study lacks sufficient power to robustly analyze sex-specific effects, such as differences in cell class proportions or the influence of sex on gene expression regulation.

### Tissue dissociation

Nuclei isolation from fresh frozen tissue was conducted following the protocol outlined by the Allen Institute for Brain Science (https://www.protocols.io/view/isolation-of-nuclei-from-adult-human-brain-tissue-eq2lyd1nqlx9/v2), with specific guidelines. All procedures were performed at 4°C to maintain sample integrity. Tissue samples (100–150 mg) were thawed on ice and homogenized in 2 mL of chilled, nuclease-free homogenization buffer. The buffer composition included 10 mM Tris (pH 8), 250 mM sucrose, 25 mM KCl, 5 mM MgCl_2_, 0.1 mM DTT, protease inhibitor cocktail (1×, 50× stock in 100% ethanol, G6521, Promega), 0.2 U/μL RNasin Plus (N2615, Promega), and 0.1% Triton X-100. Homogenization was carried out using a dounce tissue grinder with loose and tight pestles (20 strokes each; 357538, Wheaton). The nuclei solution was filtered sequentially through 70 μm and 30 μm strainers. Tubes and strainers were rinsed with an additional homogenization buffer to achieve a final volume of 6 mL. The filtered solution was centrifuged at 900 rcf for 10 minutes, after which the supernatant was discarded, leaving 50 μL of buffer above the pellet. The pellet was resuspended in 200 μL homogenization buffer, yielding a final volume of 250 μL. Next, the suspension was mixed 1:1 with 50% iodixanol (OptiPrep Density Gradient Medium, D1556, Sigma) prepared in 60 mM Tris (pH 8), 250 mM sucrose, 150 mM KCl, and 30 mM MgCl_2_. This mixture was carefully layered over 500 μL of 29% iodixanol in a 1.5 mL tube and centrifuged at 13,500 rcf for 20 minutes. Following centrifugation, the supernatant was gently removed to avoid disrupting the pellet. The pellet was resuspended in 50 μL of chilled, nuclease-free blocking buffer containing 1× PBS, 1% BSA, and 0.2 U/μL RNasin Plus. This suspension was transferred to a fresh tube, brought to a total volume of 500 μL with blocking buffer, and prepared for fluorescent activated cell sorting (FACS). For neuronal enrichment, 1 μL of NeuN antibody (1:500, Millimark mouse anti-NeuN PE conjugated, FCMAB317PE, Merck) was added to the samples, which were incubated on ice in the dark for 30 minutes. After centrifugation at 400 rcf for 5 minutes, the supernatant was discarded, leaving approximately 50 μL of buffer above the pellet. The pellet was resuspended in 500 μL of blocking buffer, passed through a 20 μm filter into FACS tubes, and stained with 1 μL of DAPI (0.1 mg/mL, D3571, Invitrogen) prior to sorting.

### Fluorescent-activated nuclei sorting

The nuclei suspension was maintained in darkness throughout the sorting process, which was developed using a flow cytometer (either BD FACSAria Fusion or BD FACSAria III) at 4°C. Gating was performed based on DAPI and phycoerythrin signals, separating the nuclei into two distinct populations: NeuN^+^ and NeuN^−^. Each sorted population was collected into separate tubes containing 50 μL of blocking buffer, with sorting continued until approximately 200,000 nuclei per population were obtained. Following sorting, the nuclei populations were centrifuged at 400 rcf for 4 minutes. The supernatant was carefully removed, leaving approximately 30 μL of buffer to resuspend the pellet. The samples were kept on ice to preserve their integrity for subsequent analysis.

### Library preparation

Library preparation from the sorted nuclei suspensions was carried out using the Chromium Next GEM Single Cell 3’ Reagent Kit v3.1 (PN-1000268, 10x Genomics). Nuclei populations were manually counted, and their concentrations were adjusted to a range of 200–1700 nuclei/μL. Following the manufacturer’s protocol (CG000204 Rev D, 10x Genomics), reverse transcription (RT) mix was added to the nuclei suspensions. For library preparation, samples were either loaded into separate lanes on the Chromium Next GEM Chip G (PN-1000120, 10x Genomics) for each population (targeting a recovery of 5000 nuclei per population) or mixed prior to loading, with proportions of 70% NeuN+ and 30% NeuN− nuclei (targeting a recovery of 5000 or 7000 nuclei). Subsequent cDNA synthesis and library preparation were conducted in accordance with the manufacturer’s instructions, utilizing the Single Index Kit T Set A (PN-1000213, 10x Genomics). Quality control and quantification steps within the protocol were performed using the Agilent High Sensitivity DNA Kit (5067-4626, Agilent Technologies) and the KAPA Library Quantification Kit (2700098952, Roche) to ensure the accuracy and reliability of the prepared libraries.

### Illumina sequencing

Pooled libraries were prepared by combining up to 19 samples for a target nucleus recovery of 5000 or up to 16 samples for a target recovery of 7000 nuclei. Sequencing was conducted on a NovaSeq S6000 platform using an S4–200 (v1.5) flow cell with eight lanes, employing a 28-8-0-91 read configuration. The sequencing was carried out at the National Genomics Infrastructure in Stockholm, Sweden.

### snRNA- seq data analysis

#### Pre-processing

Raw single-nuclei RNA sequencing (snRNA-seq) data were processed into count matrices using CellRanger (v3.0.0, 10x Genomics). The sequencing reads were aligned to the hg38 genome (GRCh38.p5; NCBI:GCA_000001405.20), incorporating both intronic and exonic sequences during the alignment process.

#### Quality Control

To identify potential doublets, we used Scrublet^37^ for each sample individually, conducting 100 iterations with automated threshold detection, default parameters, and a fixed random seed. Nuclei identified as doublets in more than 10% of the Scrublet runs were excluded. Quality control based on the distribution of unique molecular identifiers (UMIs) and unique genes detected per nucleus was then performed. Nuclei with fewer than 500 UMIs or 1200 genes were removed, as were those with more than 250,000 UMIs, over 15,000 genes, or exceeding 10% mitochondrial content. For the remaining nuclei, we modeled the logarithmic relationship between the number of unique genes and UMIs as a second-degree polynomial function. Nuclei exhibiting significant deviations from the polynomial fit—defined as a difference greater than 2000 between log-transformed gene counts and the predicted value based on UMI counts—were classified as outliers and excluded. Additionally, nuclei expressing high levels of marker genes corresponding to multiple cell types were removed. To assess this, a cell type score was calculated for each subtype (Oligodendrocytes, Microglia, OPCs, Neurons, Astrocytes, Vascular cells) for each nucleus, based on the mean expression of canonical marker genes. The score distributions were evaluated across the dataset, revealing bimodal patterns in all cases. These distributions were modeled as Gaussian mixtures, and a threshold was set at the mean of the lower Gaussian distribution plus four standard deviations. Nuclei exceeding the threshold for multiple cell types were classified as doublets and excluded. To further reduce contamination from neighboring regions, such as the claustrum or amygdala, nuclei expressing regional marker genes (*NEUROD2, TMEM155, CARTPT, SLC17A7*) were removed, with markers derived from the Allen Brain Atlas^38^. The number of nuclei excluded at each stage of this filtering process is summarized in Supplementary Fig. 1.

#### Oligodendrocytes detection

Count matrices were analyzed using Scanpy^39^ to facilitate clustering and cell type annotation, with a focus on identifying oligodendroglia. Principal component analysis (PCA) was performed, and a neighborhood graph was computed using the first 30 principal components (PCs). Clustering was conducted using the Louvain algorithm at a resolution of 0.2. The resulting clusters were annotated as either glial cells or neurons based on the expression profiles of canonical marker genes:

Astrocytes – *AQP4*, *ADGRV1*

Microglia – *CSF1R*, *FYB1*

Oligodendrocytes – *MBP*, *MOG*, *MAG*

OPCs – *PTPRZ1*, *PDGFRA*, *VCAN*

Vascular cells – *EBF1*, *ABCB1*, *ABCA9*

Neurons – *MEG3*

Oligodendrocytes and OPCs were selected, and genes were filtered based on their expression per cell. Genes detected in less than three cells were removed (1389 genes).

#### Oligodendroglia lineage classification

Oligodendroglia nuclei were projected onto the first 15 principal components (PCs) calculated on their 10% of most variable genes and re-clustered using the Louvain algorithm. Harmony^40^ algorithm was used to remove the batch effect and integrate the data.

The function scanpy.tl.rank_genes_groups from Scanpy^39^ was used to perform a differential expression analysis between the clusters through a Wilcoxon rank-sum test. Marker genes were selected manually from the top ranked genes (log foldchange > 0.5 and p-value < 0.05) to characterize and name each of the Oligodendroglia clusters as a different Oligodendroglia subclass. Markers for each subpopulation are represented in Supplementary Table 2.

#### Hierarchical analysis

Hierarchical clustering of the identified subpopulations was performed using the scanpy.tl.dendrogram function from the Scanpy^39^ package. The clustering was based on the Pearson correlation method to measure similarity between subpopulations, and the complete linkage method was applied to define the cluster structure. This approach allowed for the visualization of relationships between subpopulations based on their transcriptomic profiles.

#### Compositional analysis

The differences in cellular proportion composition between conditions and brain regions were analyzed using scCODA, a Bayesian model specifically designed to analyze changes in compositional data, particularly from single-nuclei RNA sequencing experiments. This method accounts for the inherent dependencies in compositional data, providing more accurate insights compared to both compositional and non-compositional alternatives. scCODA has been demonstrated to outperform other approaches in this domain, as described by Büttner *et al*^41^.

#### Module analysis

To identify specific gene modules associated with Oligodendroglia subpopulations, particularly those linked to *OPALIN* expression, we applied Hotspot^42^ to our subsetted dataset. Hotspot detects gene modules by leveraging co-expression patterns and analyzing single-nuclei transcriptomic data, enabling the identification of functionally related gene clusters. We used default parameters but restricted our analysis to the top 10,000 highly variable genes to focus on the most relevant features of the dataset.

#### Gene Set Enrichment Analysis

Gene Set Enrichment Analysis (GSEA) was performed using the Python package GSEApy (version 1.0.4). Pathways were considered significantly enriched if they exhibited a Normalized Enrichment Score (NES) greater than 0.5 and a false discovery rate (FDR) q-value below 0.05, to account for multiple hypothesis testing. The analysis was conducted using Gene Ontology (GO), Kyoto Encyclopedia of Genes and Genomes (KEGG), and Reactome databases as reference sets for pathway annotation and functional enrichment.

#### Correlation analysis (PDAO-1/PDAO-2/MOLA ratio vs MBP)

To estimate correlations between subpopulations altered in PD (PDAO-1/PDAO-2/MOLA) and MBP values we subsetted the data for samples showing values for all the variables (number of cells related to each subpopulation and MBP intensity). To be more precise, we calculated the percentage of cells in each sample related to PDAO-1/PDAO-2/MOL-A subpopulations and we only retained samples with more than 5 cells related to a subpopulation. Finally, we used spearman correlation test with the following rationale: (PDAO-1 or PDAO-2 percentages per sample)/MOL-A and MBP intensity.

#### Trajectory analysis

Trajectory analysis was conducted using two complementary methods. First, StaVia^43^ was applied to integrate spatial and temporal data for mapping Oligodendroglia differentiation. This method utilizes higher-order random walks to trace lineage pathways and identify intermediate states. We executed the function run_VIA with default settings, modifying only the knn parameter to 15 for consistency across analyses. The second method, Slingshot^44^, aligns single-cell data by identifying developmental lineages and connecting clusters to predict differentiation trajectories, effectively modeling the continuous maturation of Oligodendroglia. Slingshot was also run with default parameters.

#### Factor analysis

To investigate the genes associated with the differentiation process, we employed factor analysis using the scikit-learn Python package. Factor analysis is a statistical method that models observed variables as linear combinations of potential underlying latent variables, known as factors. This technique aims to identify latent structures that explain the observed variability in the dataset while minimizing noise.

In this study, factor analysis was applied to pseudotime-marked gene expression data to uncover the primary latent factors driving gene expression changes during differentiation. The input data were normalized and scaled prior to analysis to ensure consistency and comparability across genes. By leveraging factor analysis, we identified key gene sets that contribute to distinct phases of the differentiation trajectory.

#### Communication analysis

With the aim of understanding the kind of communications and the quantity of them in Oligodendroglia subpopulations, we used CellChat^45^. This method identifies ligand-receptor interactions across different subpopulations, constructing communication networks that reveal key signaling pathways involved in Oligodendrocytes development. This analysis was developed in R, and we subsetted some populations, limiting the number of cells to 10,000 (comprising 5000 Control and 5000 PD randomly selected) in the case the number of cells of the subpopulation exceeded this number.

### Immunohistochemistry with peroxidase-DAB method

Eight-μm-thick formalin-fixed paraffin embedded sections from the striatum (contralateral to samples used for snRNA-seq and spatial transcriptomics) at the level of the nucleus accumbens were immunostained in a Leica BOND RX fully automated research stainer using the BOND kit (Leica Biosystems, DS9800). Primary antibodies and concentrations were as follows: mouse anti-α-syn clone LB509 ((Thermo Fisher Scientific Cat# 18-0215, RRID:AB_2925241, 1:200), mouse anti-myelin basic protein (MBP) amino acids 129-138 clone 1 (Millipore Cat# MAB382, RRID:AB_94971, 1:250), and rabbit anti-tyrosine hydroxylase (TH) polyclonal (Cell Signaling Technology Cat# 2792, RRID:AB_2303165, 1:100). The immunohistochemistry protocol included bake and dewax steps, a 20-min-long heat-induced epitope retrieval (with ER2 solution for α-syn and with ER1 solution for MBP and TH), and hematoxylin counterstaining. Sections were coverslipped with Permount mounting media and whole-slide images were obtained under the 40× objective (numerical aperture 0.95, resolution 0.17 μm/pixel) of an Olympus VS120 virtual slide scanner (Olympus, Tokyo, Japan).

### Immunohistochemistry images analysis and Quantification

All slide images were processed and analyzed using QuPath (version 0.4.3). Quantitative analysis focused on detecting intensity features related to MBP, as well as identifying the density of TH-positive projections from the SNpc to the dorsal striatum and α-synuclein aggregations (Lewy bodies/neurites, LBs/LNs).For each image, random regions of interest (ROIs) were selected within the putamen and caudate nucleus. TH-positive projections and α- synuclein aggregation detection was performed independently using QuPath’s pixel classifier tool. Five representative samples were chosen, and within each sample, the classifier was trained using 25 TH-positive and α-syn-positive annotations and 25 TH-negative or α-syn- negative annotations, resulting in a total of 125 TH-positive and 125 TH-negative annotations, and 125 α-syn-positive and 125 α-syn-negative annotations for classification.

The data obtained from MBP intensity measurements, TH projections quantification and α- syn aggregate quantifications were further analyzed using the SciPy^43^ Python library. A two-sample t-test was applied to determine the T and p-values for each brain region and condition. For MBP analysis, the mean of diaminobenzidine (DAB) intensity per pixel within the selected ROIs was used as the metric for quantification. TH differences were assessed by comparing the percentage of positive TH annotations, calculated as the area of TH-positive measurements, divided by the total area analyzed (µm^2^). Similarly, α-syn differences were quantified based on the proportion of α-syn-positive regions relative to the total area (mm^2)^ examined in Pu or CN, respectively. Additionally, Spearman’s rank correlation analysis was performed to explore relationships between various feature pairs across different regions and experimental conditions.

### Spatial transcriptomics (Xenium)

#### Spatial Transcriptomics Platform

Spatial transcriptomic analyses were conducted using the Xenium high-plex *in situ* platform (10x Genomics), which facilitates subcellular-resolution characterization of RNA within tissue sections, similarly as previously done in Garma *et al*^46^.

#### Gene Panel Design

The Xenium technology employs oligonucleotide probes designed to quantify the expression of genes in a predefined panel. For this study, a gene panel comprising 266 genes from the Xenium Human Brain Gene Expression Panel was utilized. An additional 100 genes were selected based on our single-nucleus RNA sequencing (snRNA-seq) dataset, generating a custom gene panel (Xenium Custom Gene Expression Panel 51–100 (Z3DREH), PN-1000561, 10x Genomics).

#### Experimental Workflow

Tissue blocks from human subjects (N = 4; including Pu and CN) were retrieved from −80°C storage and transferred on dry ice to a cryostat (CryoStar NX70, Thermo Scientific). Samples were mounted on the specimen holder using Tissue Tek O.C.T. Compound (4583, Sakura) and allowed to acclimate to −20°C within the cryostat chamber for 5 minutes. Tissue sections of 10 μm thickness were obtained and placed directly within the imaging region of precooled Xenium slides (12×24 mm, PN-3000941, 10x Genomics). Adherence of the sections to the slides was facilitated by briefly warming the reverse side of the slides with gentle pressure, followed by immediate refreezing on the cryobar. Tissue-mounted slides were retained in the cryostat during the sectioning process and subsequently stored at −80°C.

Downstream processing, including probe hybridization, ligation, and rolling circle amplification, was conducted at the *in situ* Sequencing Infrastructure Unit (Science for Life Laboratory, Stockholm), following the manufacturer’s protocol (CG000582 Rev E, 10x Genomics). Background fluorescence was minimized via chemical quenching. Tissue sections were imaged using the Xenium Analyser instrument (10x Genomics), which also facilitated signal decoding and data acquisition.

#### Oligodendroglia identification

Xenium experiments provide spatial positions of all decoded reads. Therefore, the first step in processing these datasets is to segment individual cells and identify its composition. In this study, cells from Xenium experiments were defined based on default nuclear Xenium segmentation. This conservative approach minimizes the possibilities of missegmentation, as illustrated by Marco Salas *et al*^47^. Segmented cells were then preprocessed using scanpy to identify oligodendroglia. Essentially, preprocessing of the spatial dataset involved filtering out low-quality cells. Cells with fewer than 40 total counts or fewer than 10 detected genes were excluded. The data were then normalized, log-transformed, and subjected to neighbor graph construction using scanpy’s preprocessing pipeline. A UMAP embedding was generated to visualize clusters, and Leiden clustering was applied with a resolution of 0.8 to identify distinct cell populations.

Oligodendrogia clusters (clusters 0, 3, and 9) were identified based on the expression of known oligodendroglia marker gene expression (MBP, OLIG1, OLIG2, PDGFRA, among others). The selected clusters were subsetted into a new AnnData object for further analysis. Marker gene expression was visualized spatially using matplotlib and scanpy to confirm the presence and distribution of oligodendrocytes.

#### Oligodendroglia cell type assignment

Cell type deconvolution was performed using the cell2location^48^ package, which infers the spatial distribution of cell types by integrating sc-RNAseq data as a reference. First, the single-cell reference dataset, generated in this study, was preprocessed to define a knowledge base of oligodendroglia cell type-specific gene expression profiles. Genes were filtered to retain those expressed in at least five cells, representing at least 3% of the total dataset, and exceeding a mean expression cutoff of 1.12.

The spatial dataset, including only pre-defined oligodendroglia cells, was filtered to retain genes that overlapped with the single-cell reference. Cell2location’s setup function was used to prepare the spatial AnnData object for model training, specifying the sample batch as a key parameter. A regression model was trained on the reference data to estimate cell-type-specific mean expression levels. The spatial data were then analyzed using the cell2location model, which was trained for 3000 epochs with a detection alpha parameter of 200 and batch size of 10,000. Posterior distributions of cell abundances were exported to generate quantitative summaries of cell type probabilities for each spatial location. The q50 quantile (median) cell type abundances were visualized and further analyzed to identify the dominant cell types in each spatial coordinate. For oligodendrocytes, additional analysis of spatial localization and confidence intervals was performed using the posterior distributions.

#### Neighborhood Enrichment Analysis

To assess the neighborhood enrichment of oligodendrocytes, spatial interaction networks were constructed for each sample in the dataset. The AnnData object containing oligodendroglia profiled with Xenium was first subsetted by sample, and spatial neighbors were computed using Squidpy’s spatial_neighbors function with a radius of 124 um. This process was repeated for all samples, and the resulting AnnData objects were concatenated into a unified dataset.

Neighborhood enrichment was calculated using Squidpy’s nhood_enrichment function, with the cell type annotation, defined using cell2location, as the cluster key. This analysis computes z-scores representing the degree of enrichment or depletion of specific cell types in the neighborhoods of other cell types. Visualization of neighborhood enrichment was performed using a heatmap with a diverging colormap (coolwarm), highlighting significant positive and negative interactions. Parameters such as color scale limits (vmax = 200, vmin = -200) and figure size were adjusted to optimize clarity and interpretability. Overall, this method provides insights into the spatial interactions and organization of oligodendrocytes in their local tissue environment, enabling a deeper understanding of their roles in tissue architecture and function.

#### Module identification using Hotspot

We used Hotspot^42^ to identify gene expression modules in oligodendroglia profiles derived from Xenium spatial transcriptomics data. Hotspot identifies informative genes and gene modules based on how gene variation aligns with cell similarity metrics. Informative genes show expression patterns concordant with local cell similarity. In summary, we initialized Hotspot with the following parameters:

- Layer key: ‘raw’ for raw expression values.
- Model: Negative binomial (danb).
- Latent obsm key: ‘X_pca’.
- UMI counts key: Total UMI counts per cell.

A k-nearest neighbors (kNN) graph (20 neighbors, unweighted) was created to compute gene autocorrelations. Genes with FDR < 0.05 were selected as significant. Local correlations for these genes were computed using parallel processing. Gene modules were defined using a minimum threshold of five genes, with core genes meeting an FDR threshold of 0.05. Module scores for each cell were calculated and stored. Differences in module scores between PD and control groups were analyzed using the Wilcoxon rank-sum test. Violin plots visualized score distributions, with test statistics and p-values annotated.

#### Niche identififcation using NicheCompass

NicheCompass^49^ was used to identify cellular domains by integrating spatial transcriptomics data across all samples. A latent graph embedding was computed using a graph convolutional network (GCN) model with categorical covariates such as replicates (Code availability). NicheCompass was run for 400 epochs with regularization to optimize ligand-receptor-based niche identification. Clustering was performed on the latent embeddings using the Leiden algorithm with a resolution of 0.2. After clustering, niches with fewer than 200 cells were filtered out, leaving three robust niches, that could be annotated as white matter (WM), gray matter (GM), and vascular niches based on their expression and cellular compostion. These niches were characterized by their communication program activity, spatial localization, and enriched gene targets.

### Statistical analysis

Statistical analyses were conducted using Python-based computational tools. Differential expression analysis was performed using the Wilcoxon rank-sum test, with marker genes selected based on a log fold change >0.5 and p-value <0.05. Hierarchical clustering of subpopulations was conducted to assess similarity based on Pearson correlation. Cellular compositional differences between conditions and brain regions were analyzed using a Bayesian model designed for sn-RNAseq data. GSEA statistical significance was assessed using permutation-based methods (a Kolmogorov-Smirnov-like statistic), which compares the observed enrichment of genes in a gene set against a random distribution. Spearman’s rank correlation was used to examine relationships between specific features, such as the percentages of PDAO-1 or PDAO-2 and MOL-A/MBP intensity. Two-sample t-tests were used to compare MBP intensity, TH positive dots, and α-syn-positive areas across conditions and brain regions, generating T and p-values. Sample sizes were determined based on prior studies and experimental feasibility. Detailed statistical results and methodologies are presented in the figures and legends.

## Results

To elucidate the genetic networks and underlying molecular changes in OLs lineage cells within the context of PD in the dorsal striatum, we isolated and sequenced individual nuclei from freshly preserved human brain tissue samples obtained from CN and Pu of N = 27 Controls and N = 36 PD donors. By employing snRNA-seq through a well-established workflow (Fig. 1A), we obtained 200,000 individual nuclei associated with OLs lineage cells (Fig. 1C). Notably, this dataset represents the most extensive compilation of human OLs lineage cell transcriptomic information from the dorsal striatum available to date. Control and PD groups were matched by age but not by sex due to the higher prevalence of PD in men (Fig. 1C). The average number of cells per donor and striatum (including CN and Pu) was approximately 3000 (Fig. 1C). To characterize and confirm the loss of dopaminergic neuron projections and the presence of LBs/LNs in PD samples, we performed immunohistochemical analysis for tyrosine hydroxylase (TH) and α-synuclein (α-syn) in the contralateral CN and Pu of MADRC Control (N = 10 (α-syn), N = 8 (TH)) and PD (N = 8 (α-syn and TH)) donors, (Fig. 1B, Supplementary Fig. 2), for whom formalin-fixed paraffin-embedded sections were available. The analysis of immunoreactivity revealed a significant reduction in TH immunoreactivity (Control = 1.66 +- 0.58 (SEM) versus PD = 0.21 +- 0.05 (SEM), p = 0.027, t = 2.47) and a marked increase in α-syn immunoreactivity (Control = 0.004 +- 0.0017 versus PD = 0.01 +- 0.002, p=0.035, t = 2.31) specifically in the Pu of PD patients (Fig. 1B, Supplementary Fig. 2), reflecting the expected loss of TH-positive dopaminergic neuron projections and the aggregation of α-syn in LB/LNs in PD, respectively.

**Figure 1.**
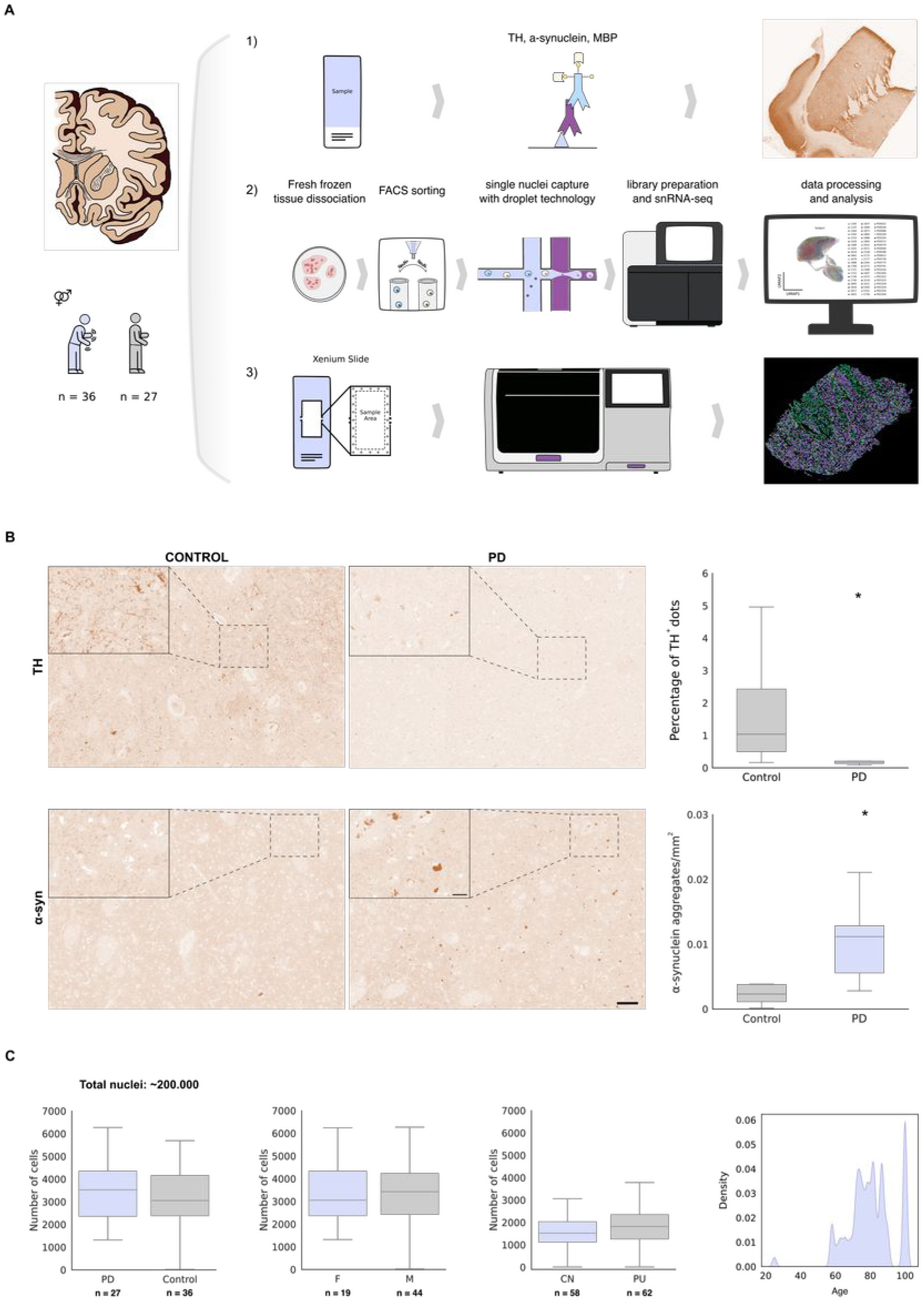
The human dorsal striatum shows great diversity of oligodendroglia and specific associated subclasses for PD. **A.** Schematic overview of the experimental design. **B.** Immunohistochemical analysis of Tyrosine Hydroxylase (TH) and α-synuclein (α-syn) in the putamen. Box plots illustrate the distribution of marker expression in Control and Parkinson’s disease (PD) conditions. Statistical significance was determined using two sided t-tests, with significance indicated with an asterisk (TH analysis: p = 0.027, t = 2.47; α-syn analysis: p = 0.03, t = 2.39). Scales are set at 100 μm for low power images and at 20 μm for insets. **C.** Quantitative analysis of sequenced nuclei for OLs and OPCs, categorized by condition, sex, and region, along with the age distribution frequencies of the subjects.

After implementing a robust quality control process (Supplementary Fig. 1), we conducted an initial clustering to identify four main classes: Oligodendrocyte Precursor Cells (OPCs) expressed *VCAN* as marker gene; Committed OPCs (COPs) expressed *GPR17* as marker gene; and Mature Oligodendrocytes (MOL) were identified by the expression of *MBP*, and further subdivided based on the expression levels of *OPALIN* or *SLC5A11* marker genes (Fig. 2. B). These classes were represented in both control and PD as well as in CN and Pu (Fig. 2A, B, Supplementary Fig. 3). Subsequently, cell classes were subclustered identifying 15 distinct subpopulations. Due to their clear enrichment in PD, we annotated some of these as PD-associated (PDA) (Fig. 2B, D). OPCs displayed four subclasses: OPCs-A (expressing the gene markers *VCAN/GPC6/SGCZ*), PDAOPCs (*VCAN/MT3/CD81*), OPCs-C (*VCAN/SEMA3E/GRIA4*), and OPCs-D (*VCAN/ST18*). MOLs expressing *OPALIN* comprised five subpopulations: MOL-A (*LAMA2/KIF6*), MOL-B (*FTH1/CRYAB/DMD*), PDAO-1 (*DBNDD2/FTL/HSPA1A*), PDAO-2 (*CTNNA2/ARHGAP24*), and MOL-F (*LURAP1L-AS1, LINC00609, LINC01608*). Within the *SLC5A11^+^* population, distinctions were made between PDAO-3 (*ACTN2*/*MT-ND3*) and MOL-G (*LINC01505, PLEKHG1, SLC25A29*) (Fig. 2B, C).

**Figure 2.**
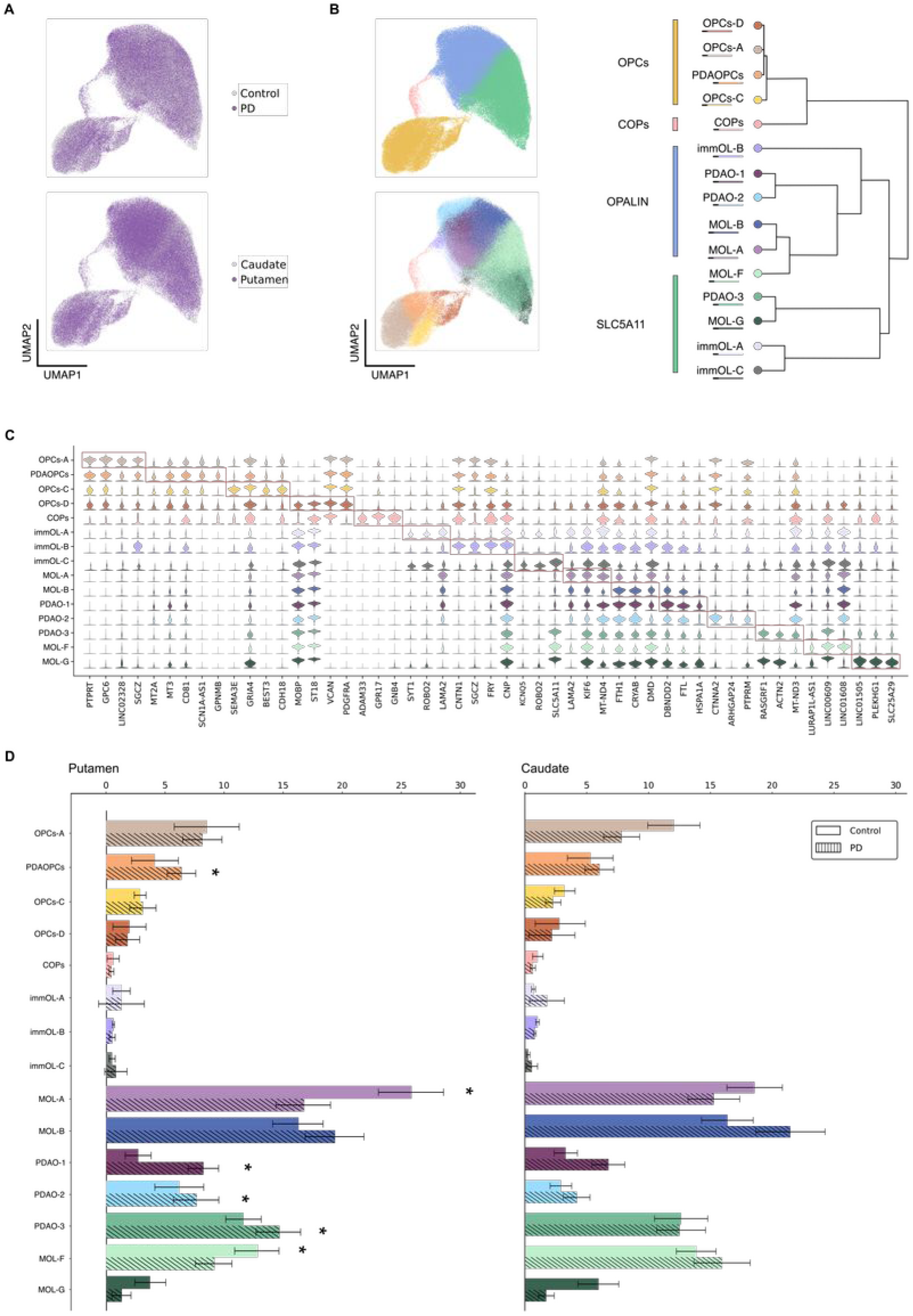
PD-associated clusters show specific transcriptomic signatures including stress-reactive and immunological responses. **A.** UMAPs display the uniform distribution of cells from each region and condition across all subpopulations. **B.** (**Top left**) UMAP representing principal oligodendrocyte (OL) and oligodendrocyte progenitor cells (OPC) classes: OPCs, COPs, MOLs-*OPALIN*^+^, MOLs-*SLC5A11*^+^. (**Bottom left**) UMAP visualization depicting OLs and OPCs subpopulations, with distribution within subclasses based on transcriptomic similarity, as illustrated in the dendrogram (**Right**). **C.** Stacked violin plots representing marker expression profiles for each subpopulation. **D.** Bar plots showing the percentages of each subpopulation, calculated based on the total number of nuclei in each condition. Asterisks (*) indicate significantly altered populations based on scCODA p-value of confidence (< 0.2).

The identified subpopulations represent a continuum; therefore, they do not correspond to distinct OL subtypes, but rather reflect different states within a specific subpopulation, exemplified by either different stages of differentiation or activity specialization. This is particularly evident in MOLs, where most subpopulations share common markers, with their key distinction lying in their gene expression levels rather than in the presence or absence of specific markers. These findings suggest that transcriptomic variability within MOLs is mainly driven by different functional states rather than by the emergence of discrete cell types.

After identifying distinct subpopulations within the OLs lineage, we characterized each by differential gene expression analysis (Fig. 2C, Supplementary Table 2). Of note, the disease-specific clusters (PDAOPCs, PDAO-1, PDAO-2, and PDAO-3) exhibited upregulated expression of heat shock proteins (HSP) families 70 and 90, including *HSPA1A, HSPB1, HSP90AB1, HSPA1B, HSPA8, HSP90AA1*, and *HSPA5*, as well as *CRYAB* and *DNAJ* family members. Thus, the elevated levels of these chaperone genes in PDAO-1 and PDAO-2 subpopulations underscores an enhanced cellular response to stress and their involvement in protein folding and proteostasis processes within the OLs lineage (Fig. 2C).

To assess the potential disease and region specificity of these subclasses, we conducted a compositional proportion analysis using scCODA. In Pu, PD samples exhibited modest increases in PDAO-1 (+5.6%), PDAO-2 (+1.5%), PDAO-3 (+3.0%), and PDAOPCs (+2.3%), alongside a reduction in MOL-A (-9.1%) and MOL-F (-3.7%) (Fig. 2D). Notably, these PD-induced alterations were only significant in Pu, although the same tendency was observed in CN (Fig. 2D). Thus, in the striatum, particularly in the Pu, a subset of OLs exhibit a prominent stress response in PD (Fig. 2D). Furthermore, to control for potential sex-related effects, we performed a similar compositional proportion analysis stratified by sex and found no significant differences between males and females in any of the identified subpopulations, suggesting that the observed changes are independent of sex (Supplementary Fig. 4).

We next focused on *OPALIN^+^* MOLs as they showed to be the most affected class with three altered populations in PD (Fig. 2B, D). To confirm the gene markers associated with the *OPALIN^+^* subpopulations, including PDAO-1 and PDAO-2, and identify gene co-expression patterns and cellular programs associated with those gene markers, we performed Hotspot analysis. This analysis revealed several gene modules, of which Modules 4 (genes *KCNIP4*, *RBFOX1*) and 5 (*MDGA2*, *CNTN1*, *LUZP2*, *FRY*, *GRIK2*) were associated with immature OLs-A (immOL-A) and immature OLs-B (immOL-B), respectively. Interestingly, Modules 10 and 14, increased in MOL-A, included mitochondrial genes and genes regulating projection development (*GRID2*, *ZFPM2*), respectively. PDAO-1 showed increased expression of Modules 1 and 2, which were associated with HSP genes (*CRYAB*, *FTH1*, *FTL*, *HSPA1A*, *HSPA1B*, *HSPH1*), while PDAO-2 was linked to Modules 6 and 13, which are enriched in response to cytokines and antigen processing and presentation (*ETV5*, *QDPR*, *ARHGAP24*, *CTNNA2*) (Supplementary Fig. 5).

In light of the significant changes observed within the mature OLs associated with PD (included in the *OPALIN^+^* and *SLC5A11^+^*subclasses), we conducted a Gene Set Enrichment Analysis (GSEA) on the most affected subpopulations, employing established databases such as Gene Ontology, KEGG, and Reactome, to better understand the functional pathways implicated in PD. Our analysis focused on four key comparisons based on the compositional and dendrogram analyses: MOL-A vs PDAO-1 and 2, PDAO-3 vs MOL-F, and PDAOPCs vs OPCs-A, based on *OPALIN*, *SLC5A11*, and *PDGFRA* expression levels.

The MOL-A vs PDAO-1 and 2 contrast revealed a noteworthy pattern of enrichment within the MOL-A subtype (decreased in PD in Pu), with substantial involvement in essential myelination biological processes, including Cell-Cell and Focal Adhesion, Axon Guidance, Synapse Organization, Positive Regulation of Cell Growth and GTPase Activity, Regulation of Microtubule-based Process, CDC42 GTPase Cycle, RAC1 GTPase Cycle, Regulation of lipid metabolism, Netrin-1 signaling and RHOA GTPase Cycle, ERBB signaling pathway (Fig. 3A, Supplementary Table 3). On the other hand, PDAO-1 (increased in PD in Pu) showed enrichment in ATP Synthesis Coupled Electron Transport, Response to Unfolded Protein, Chaperone Cofactor-Dependent Protein Refolding, Oxidative Phosphorylation, Signaling by ROBO receptors, Antigen processing-cross presentation, Chaperone-mediated autophagy (Fig. 3A, Supplementary Table 3). Also, PDAO-2 demonstrated an immunological profile, with processes Antigen processing-cross presentation, Regulation of NF-kB signaling, Downstream signaling events of B cell receptor, and TNFR2 non-canonical NF-kB pathway, besides HSF1 activation and Chaperone-mediated Autophagy (Fig. 3A).

**Figure 3.**
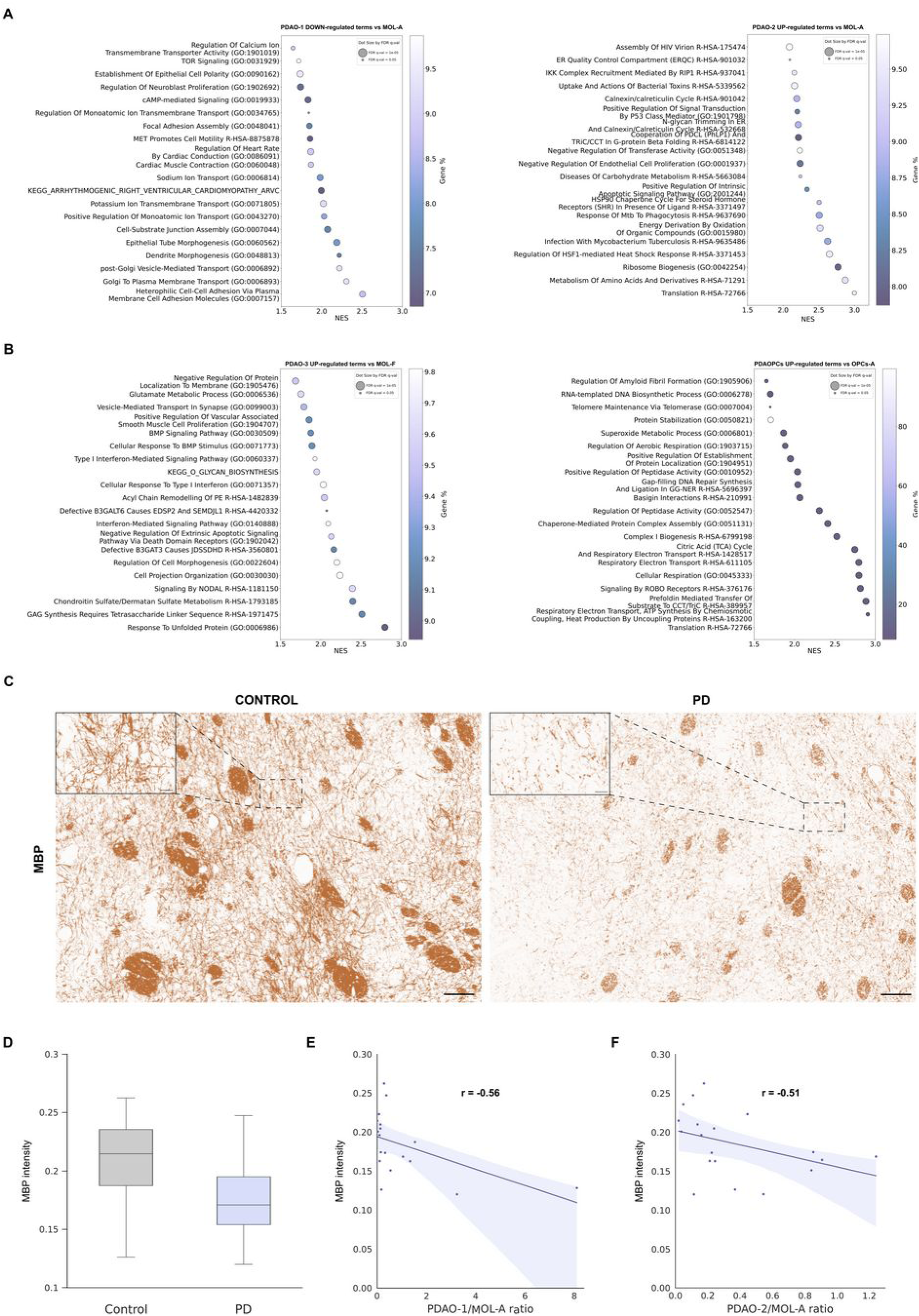
Disrupted Myelination and Key Pathways in Parkinson’s Disease-Associated Oligodendrocyte Subpopulations. **A. (Left)** GSEA results highlighting the top 20 enriched terms (from GO, KEGG, and Reactome databases) with reduced expression in PDAO-1 compared to MOL-A, ranked by gene ratio. **(Right)** Top 20 enriched terms with increased expression in PDAO-2 relative to MOL-A, also ranked by gene ratio. **B. (Left)** Top 20 enriched terms showing upregulation in PDAO-3 compared to MOL-F, based on gene ratio. **(Right)** Upregulated terms (top 20) in PDAOPCs relative to OPCs-A, ranked by gene ratio. **C.** Immunohistochemical analysis of MBP in the Pu region under Control and PD conditions. Scales are set at 100 μm for the low power images and at 20 μm for their insets **D.** Violin plots illustrating the distribution of MBP expression in Control and PD conditions. p-value = 0.047, t = 2.11. **E.** Scatter plot depicting the correlation between the ratio of PDAO-1 to MOL-A cells and MBP expression. r = -0.56, p-value = 0.02. **F.** Scatter plot showing the correlation between the ratio of PDAO-2 to MOL-A cells and MBP expression. r = -0.51, p-value = 0.02.

The second comparative analysis, PDAO-3 vs MOL-F, revealed several key pathway alterations related to myelination and myelin integrity. In PDAO-3, pathways related to positive regulation of autophagy, long-chain fatty acid metabolic processes, sphingolipid metabolism, and response to cytokines and unfolded protein response were significantly upregulated (Fig. 3B, Supplementary Table 3). Conversely, processes critical for myelin assembly, including regulation of cell projection organization, synapse assembly, Rho protein signal transduction, integrin signaling, and the Ephrin receptor signaling pathway were notably downregulated in PDAO-3, indicating a decline in these processes in the context of PD (Supplementary Fig. 6, Supplementary Table 3).

Finally, the OPCs-A vs PDAOPCs comparison demonstrated an upregulation of RAC1 GTPase cycle, CDC42 GTPase cycle, RHOA GTPase cycle, RAC3 GTPase cycle and Cell-Cell communication in OPCs-A (Supplementary Fig. 6, Supplementary Table 3), whereas PDAOPCs were enriched in processes as Citric acid (TCA) cycle and respiratory electron transport, PINK1-PRKN mediated mitophagy, antigen processing-cross presentation, binding and uptake of ligands by scavenger receptors (Fig. 3B, Supplementary Table 3). These results suggest an upregulation of energy metabolism and phagocytosis in PDAOPCs.

Furthermore, to assess the impact of PD on myelin quantity, we conducted a quantitative immunohistochemistry analysis using the Myelin Basic Protein (MBP) marker, which is indicative of myelination levels (Fig. 3C). The results revealed a decrease in MBP immunoreactivity within the Pu in PD vs Control donors (Control = 0.21 +- 0.01, PD = 0.17 +- 0.009, p-value = 0.047, t = 2.11) (Fig. 3D). In addition, to further elucidate the relationship between MBP immunoreactivity in the Pu and the relative abundance of OLs subpopulations, we conducted a correlation analysis between MBP immunoreactive signal and the ratios of PDAO-1 and PDAO-2 to MOL-A, reflecting the aforementioned shifts in cell population proportions in PD. We observed a significant negative correlation, indicating that as the ratio of PDAO-1/MOL-A (r = -0.56, p-value = 0.02) and/or PDAO-2/MOL-A (r = -0.51, p-value = 0.02) increases (suggesting a shift of MOL-A to PDAO-1 and/or PDAO-2), MBP levels decrease (Fig. 3E, F).

To understand the normal differentiation process and identify deviations from normality associated with PDAOs clusters in PD, we conducted trajectory inference analyses using the VIA and Slingshot algorithms. To infer the normal differentiation pathway, we considered all OL clusters except PDAOs, with samples from both conditions and regions. The differentiation process we identified is as follows: precursor cells, specifically OPCs-A, represent the original state and undergo differentiation into OPCs-C. These then further mature into COPs and immOL-B. Subsequent differentiation toward MOLs progresses through MOL-A and MOL-B populations, which are implicated in the initiation of myelination, as evidenced by the expression of Laminin-2 (*LAMA2*) and other cell adhesion molecules (CAMs). As differentiation advances, the MOL-F population emerges, characterized by the expression of genes associated with cholesterol metabolism, a process critical for the myelin sheath wrapping characteristic of mature oligodendrocytes (Fig. 4A). Intriguingly, we identified a previously unreported branch in the differentiation trajectory (Supplementary Fig. 7): OPC-D transitions through immOL-C to immOL-A, ultimately converging toward MOL-A/MOL-F. Moreover, immOL-C is also linked to MOL-E, suggesting a potential alternative pathway facilitating direct differentiation and maturation processes (Fig. 4A).

**Figure 4.**
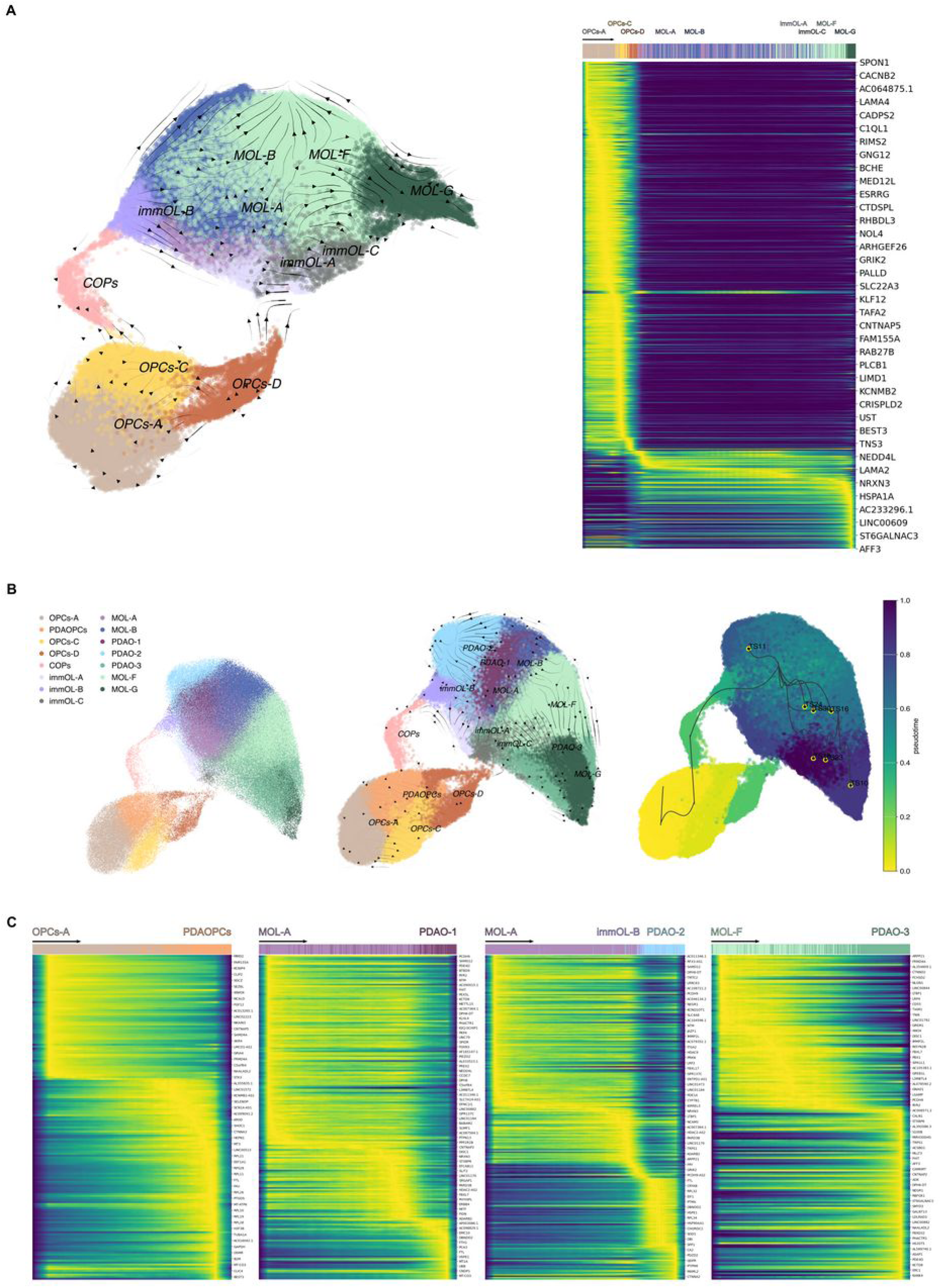
Trajectory Inference Reveals Disrupted Oligodendrocyte Differentiation Pathways in Parkinson’s Disease. **A. (Left)** Pseudotime UMAP generated via StaVia analysis, excluding the expanded populations associated with PD. Arrows depict the progression of the differentiation trajectory, which proceeds from oligodendrocyte progenitor cells (OPCs) through COPs/OPCs-D, followed by immature oligodendrocytes (immOLs), and culminating in the transition from MOL-A to MOL-G. **(Right)** Heatmap illustrating the expression of key genes involved in this differentiation process. **B.** UMAP visualizations of differentiation trajectories in PD-associated clusters. Pseudotime analysis highlights a convergence point at MOL-A, from which the differentiation diverges into PDAO-1 and PDAO-2 clusters. **C.** Schematic representation of genes associated with differentiation pathways, tracing potential origins and pathways leading to the development of PD-associated clusters.

When analyzing all the clusters, including those PDAOs, and samples from both conditions and regions, we observed deviations in this normal differentiation trajectory. PDA clusters appeared to originate from specific healthy populations, exhibiting altered differentiation pathways in the diseased striatum. Specifically, PDAO-1 likely emerged directly from MOL- A in PD striatum, reflecting a disruption in the normal differentiation pathway of MOL-A to MOL-B and MOL-F. PDAO-2 showed a more ambiguous origin, potentially involving disruptions at the MOL-A and/or immOL-B differentiation stages, indicative of a complex pathological process (Fig. 4C). In the case of PDAO-3, it appeared to arise from perturbations in MOL-F, suggesting PD-associated alterations in this late-stage differentiation population. Finally, the same applied to PDAOPCs, which originated from OPCs-A, highlighting disruptions in the earliest stages of oligodendrocyte lineage differentiation.

To further investigate the molecular mechanisms underlying these differentiation pathways, we applied factor analysis to pseudotime-ordered gene expression data, which enabled us to identify distinct genetic signatures underpinning the differentiation trajectories (Fig. 4C, Supplementary Table 4), highlighting potential molecular mechanisms involved in disease-associated alterations. Factor analysis revealed specific genetic signatures involved in the transitions observed in normal differentiation, including genes critical for myelination, cholesterol metabolism, and cell adhesion (*LAMA4, LAMA2, MOBP, ERBB4*). When applied to pathological trajectories, factor analysis highlighted specific gene dysregulations associated with PDA clusters, providing insights into molecular mechanisms disrupted by disease (Fig. 4C). These findings underscore the role of aberrant gene regulation in driving pathological deviations from normal oligodendrocyte differentiation.

We further aimed to examine potential intercellular communication among the identified subpopulations and to determine whether communication patterns differ between Control and PD striatum. With CellChat, we evaluated outgoing and incoming signaling dynamics based on the expression of known ligand-receptor pairs across subpopulations. Our results revealed significant disruptions in communication patterns within PD-associated clusters. Specifically, the analysis demonstrated a marked reduction in the expression of extracellular matrix (ECM) and adhesion molecules genes, such as *NCAM, CADM*, Laminin, and *CDH1*, in PDAO-1 compared to MOL-A, the latter exhibiting the highest expression levels of these molecules within the *OPALIN^+^* class (Fig. 5A, B). These findings suggest a weakening of adhesion and ECM-mediated signaling in PDAOs subpopulations. Indeed, there was a sharp decline in outgoing signals linked to myelination, particularly those associated with Laminin (*LAMA2*), in both PDAO-1 and PDAO-2. Furthermore, critical cell adhesion molecules (e.g., *NCAM, CADM,* and *CDH1*) were nearly absent in PDAO-1, highlighting substantial deficits in the ability of this cluster to engage in adhesion-based communication, likely contributing to the broader dysregulation of cellular networks in PD (Fig. 5B).

**Figure 5.**
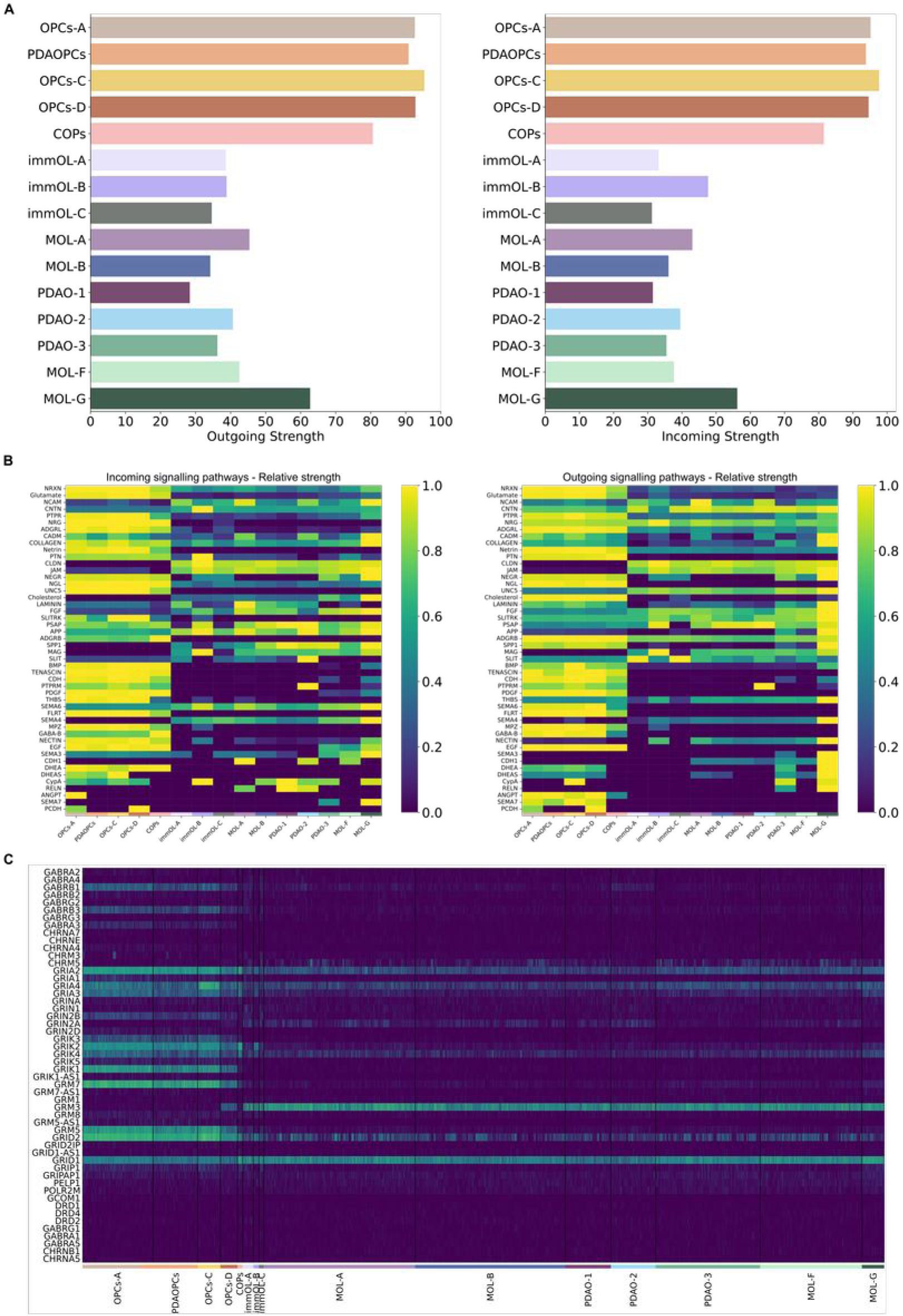
Altered intercellular interactions and neurotransmitter signatures in Parkinson’s disease-associated oligodendrocyte subclasses. **A.** Bar plots illustrating the outgoing and incoming communication strengths for each subpopulation, as determined by ligand-receptor interaction analysis. **B.** Relative signaling pathway strengths are depicted for each subpopulation, highlighting distinct communication profiles. **C.** Heatmap displaying the expression levels of neurotransmitter-associated markers identified in at least 50 cells within any given population, showcasing distinct molecular signatures across subpopulations.

In contrast, cholesterol-related molecules, particularly within the *SLC5A11^+^* class, were elevated in these subpopulations (PDAO-3, MOL-G), aligning with the distinct biological profile of these populations demonstrated with hierarchical clustering, annotation, and GSEA analyses. Notably, the PDAOPCs subpopulation exhibited the lowest levels of general communication molecules among precursor cells, further suggesting disrupted intercellular interactions in early-stage oligodendrocyte precursors in PD (Fig. 5B).

Another striking finding from the CellChat analysis was the distinct upregulation of glutamate-related signaling molecules in OPC subpopulations, particularly in PDAOPCs. This observation prompted a deeper investigation into neurotransmitter signaling profiles across oligodendrocyte subpopulations (Fig. 5C). While the most pronounced differences were observed between OPCs and MOLs, key neurotransmitter-associated genes were also found to be differentially expressed in PD-specific clusters. For instance, *CHRNA10* and *GABRG1* were highly expressed in PDAO-2, while *GABRR2* was significantly upregulated in both PDAO-1 and PDAO-2. These findings point to a distinct alteration in neurotransmitter signaling in PD-associated oligodendrocytes, likely contributing to their functional deficits.

To validate the cell populations identified through snRNA-seq and further analyze other tissue-related variables, we employed spatial transcriptomics. This technique preserves the spatial organization of gene expression within the tissue, enabling the direct mapping of snRNA-seq-defined cell populations to their native anatomical contexts. By integrating these high-dimensional data modalities, we evaluated the robustness and biological relevance of snRNA-seq-derived clusters, ensuring that the identified populations are spatially resolved and assessing whether they localize to distinct striatal regions or whether they correspond to distinct functional domains within the same region.

Through spatial transcriptomics, we profiled 435,157 oligodendroglial cells. Using the cell2location model, we transferred snRNA-seq annotations to cells identified as oligodendrocytes or OPCs in the spatial dataset (Fig. 6A). Analysis of the 366 profiled genes enabled the identification of all snRNA-seq-derived subpopulations within the oligodendrocyte lineage across both CN and Pu in the spatial transcriptomic dataset (Fig. 6A).

**Figure 6.**
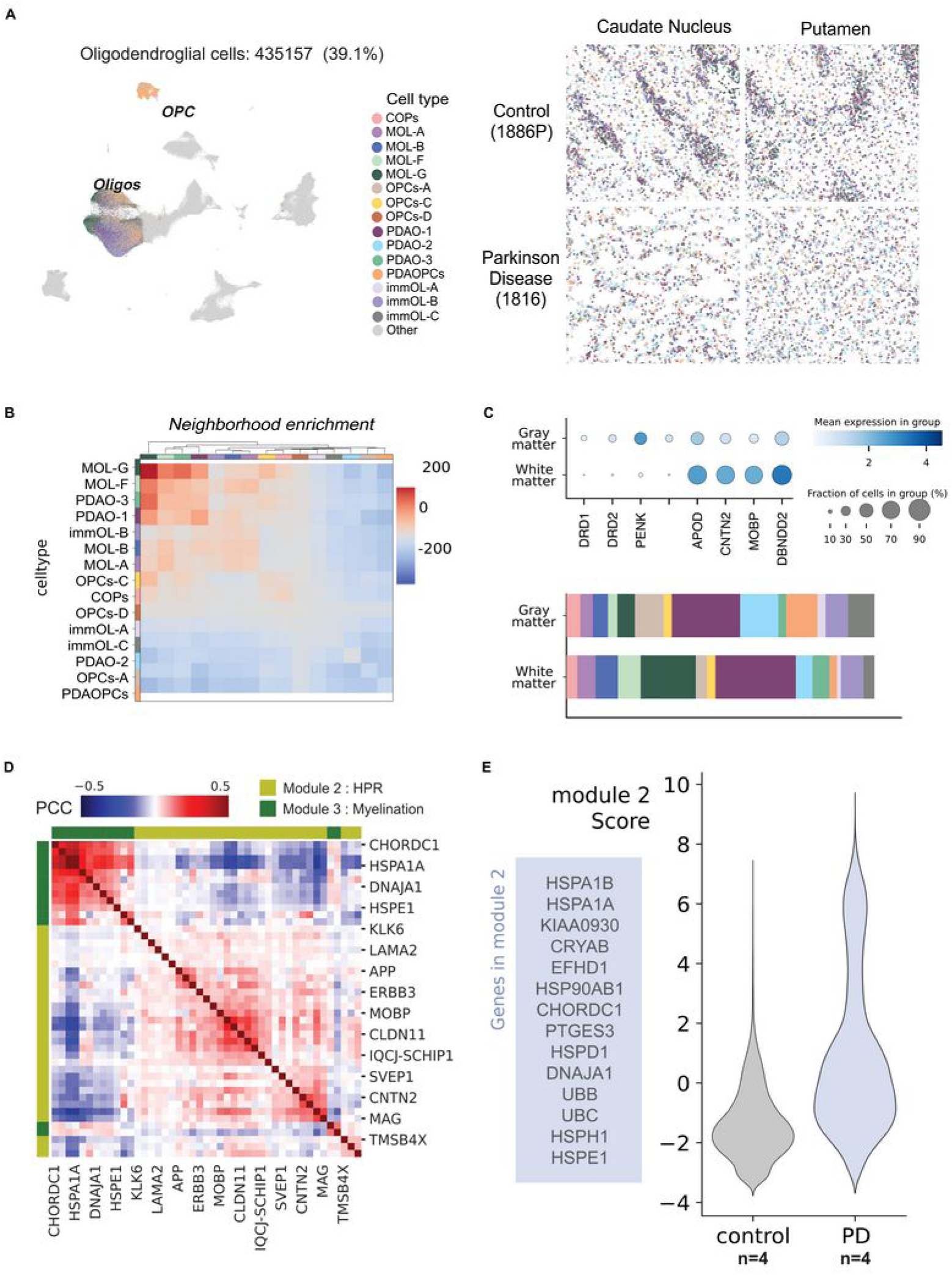
Spatial Mapping Reveals Oligodendrocyte Subpopulations and Myelin-Related Alterations in Parkinson’s Disease. **A. (Left)** UMAP illustrating oligodendrocyte populations profiled through spatial transcriptomics, corresponding to the populations identified via snRNA-seq. **(Right)** Spatial distribution of oligodendrocyte subpopulations across CN and Pu. **B.** Heatmap depicting neighborhood enrichment patterns for all identified subpopulations, highlighting spatial colocalization. **C.** Spatial localization of oligodendrocyte subpopulations within the gray matter (GM) and white matter (WM) areas. These regions were categorized based on marker expression (**Top**). Bar plots illustrate the distribution of oligodendroglial subpopulations across both GM and WM **(Bottom)**. **D.** Heatmap represents the distribution of module 2 (Heat Shock Protein Response, HSPR) and module 3 (myelination), demonstrating a negative correlation between these modules. **E.** Violin plots showing an increased expression of HSPR module in PD.

A noteworthy observation was the spatial distribution of these subpopulations within the tissue. Neighborhood enrichment analysis identified four main colocalizing subpopulations: PDAO-1, PDAO-3, MOL-F, and MOL-G (Fig. 6B). PDAO-3, MOL-F, and MOL-G were characterized by their maintenance of a myelin-related profile and appeared to be predominantly located within the white matter tracts that cross through the striatum (Fig. 6C).

To better understand the molecular changes specific to PD, we conducted a hotspot analysis to identify distinct gene expression modules that show significant variance between Control and PD donors. This analysis highlighted two key modules with notable differences: Heat Shock Protein Response (HSPR) and Myelination. Specifically, we observed a significant upregulation of the HSPR module in PD, accompanied by a marked downregulation of the Myelination module, pointing to opposing trends in these two pathways (Fig. 6D, 6E).

The HSPR module, associated with cellular stress responses, exhibited strong correlations among genes encoding heat shock proteins, including *HSPA1A, DNAJA1, HSP90AB1*, and *CRYAB*. This module showed elevated activity in PD, suggesting an amplified cellular stress response in affected cells. Importantly, the activity of the HSPR module in PD was strongly correlated with the transcriptional signatures of PDAOs, further reinforcing the robustness of this finding (Fig. 6D, 6E).

In contrast, the Myelination module, which encompasses genes critical for myelin formation and maintenance, such as *MAG, MOBP,* and *CNTN2*, was significantly downregulated in PD, suggesting an impairment of myelination processes. The reduced co-expression among genes in this module further underscores the disruption of myelin-related pathways in PD (Fig. 6D, 6E).

Interestingly, the activity of the HSPR and Myelination modules appeared to be inversely related. In PD samples, the increased activity of the HSPR module coincided with a decrease in the Myelination module, suggesting a potential trade-off between cellular stress response mechanisms and the maintenance of myelin. This interplay likely reflects a previously unrecognized critical aspect of PD pathophysiology, where oligodendrocyte energy and/or machinery resources are likely diverted to manage stress, compromising other vital processes like myelination (Fig. 6D).

## Discussion

PD is a complex neurodegenerative disorder with no cure and unknown trigger in the vast majority of sporadic cases. Understanding the different key alterations with cell type specificity is crucial to develop treatments that can halt the progression of the disease. Traditionally PD has been associated with dopaminergic neuron degeneration and Lewy bodies. Recently, oligodendroglia has emerged as a potential critical cell type in this disease^10,32,50,51^, but its involvement has not been investigated in depth. Characterizing the diversity of OLs and their precursors, as well as how they are affected in one of the most impacted brain regions in PD (the dorsal striatum), is essential for uncovering the mechanisms underlying oligodendroglial dysfunction and its contribution to neurodegeneration.

In this work we have built the largest dataset, to our knowledge, of snRNAseq from CN and Pu samples from PD and Control donors and applied one of the most recent and sensitive spatial transcriptomics approaches in order to delineate the main transcriptomic changes of oligodendroglia with their neuroanatomical correlation in this neurodegenerative disorder.

The complexity of OLs and OPCs has only recently been elucidated in specific brain regions^20,31,52–54^. In our study we analyzed more than 200K nuclei from CN and Pu of 63 donors identifying 4 main classes and 15 distinct subclasses. The main classes (OPCs, COPs, *OPALIN*^+^ MOLs and *SCL5A11^+^*) ^10,13,20,31,53–55^.

Among our subclasses, MOL-A and G (Int0 and Int3 in Sadick *et al*^13^), and immOL-B (Int4 in Sadick *et al*^13^ and FRY population in Smajic *et al*^10^) have been previously identified. Importantly, our large study enabled the identification of novel subclasses, including OPCs- C, OPCs-D, MOL-F, and immOL-A and C, as well as those associated with the disease, such as PDAOPCs and PDAO-1, PDAO-2, and PDAO-3. While MOL-B and PDAO-2 showed similarities with previous described populations, they have a distinct transcriptomic profile (Supplementary Table 5): PDAO-2 share similarities with the disease-associated OLs (DAOs) identified in MS by Falcão *et al*^56^, including pathways involved in antigen processing and presentation via major histocompatibility complex class I and II (MHC-I and -II), and immune OLs (imOLs) from Jäkel *et al*^19^ (related to OPC1 in Falcão *et al*^56^) sharing *ARHGAP24* and *CTNNA2* among others. Although DAOs have been observed in neurodegenerative diseases like MS (Falcão *et al*^56^) and AD (Jäkel *et al*^19^), the transcriptomic profile of PDAO-2 demonstrates unique characteristics (Supplementary Table 5), suggesting a disease-specific role in PD.

Overall, the transcriptomic changes were more prominent in Pu than CN, as illustrated by changes in OL composition (Figure 2D), which is consistent with the more severe reduction in TH and MBP immunoreactivity in Pu versus CN and with the well-established spatio- temporal progression of the nigrostriatal pathway degeneration in PD ^57,58^.

HSPs alterations have been linked with microglia in the midbrain in PD^10^, and with specific signatures of OLs in PD^10,59^, but its role at the population level in OLs has not been described before. The elevated expression of HSP-related genes, including *HSPA1A*, *HSPB1*, and *HSP90AB1*, alongside increased chaperone-associated gene levels, highlights its potential role in stress adaptation and prevention of protein aggregation^60^. Intriguingly, α-syn aggregation in oligodendrocytes is the neuropathological hallmark of multiple system atrophy (MSA) but is not a feature of PD. Trajectory inference analysis indicates that PDAO-1 likely derives from the MOL-A lineage and it exhibits significant impairments in myelination-related processes compared to MOL-A, suggesting a maladaptive response in OLs during PD progression. In addition, PDAO-1 displays reduced expression of key molecules involved in myelin assembly (actin cytoskeleton-related genes and cell adhesion molecules like laminins and cadherins) and intercellular signaling with neurons, pointing out a disruption in the formation of new myelin, perhaps contributing to the loss of white matter integrity in PD striatum that we observed. Demyelination and/or remyelination failure have not been established as a pathological hallmark in PD although a few works have pointed in that direction^61,62^ and support our findings.

Identifying PDAO-2 as a distinct OLs subpopulation with an immunological profile offers intriguing insights into PD pathogenesis. PDAO-2 is characterized by increased expression of neurotransmitter receptors, particularly *GABRG1* and *GABRR2*, along with markers typically associated with immune response pathways (*HLA-C, HLA-A*). This is consistent with research showing that OLs can adopt immune-like properties in neurodegenerative conditions, engaging in processes such as antigen presentation and releasing pro-inflammatory cytokines (Jäkel *et al*^19^ and Falcão *et al*^56^). In PD, OLs may influence microglia and astrocyte activity, contributing to the chronic neuroinflammatory environment^63–65^. Furthermore, reduced expression of myelination-associated communication molecules like Laminin^66^ and Opalin^67^ in PDAO-2 may indicate a shift from a supportive role in myelination toward an immunomodulatory function. This aligns with findings in other neurodegenerative diseases, such as MS and Alzheimer, where OLs exhibit impaired myelination and participate in immune signaling, exacerbating inflammation and tissue damage^68,69^. Thus, PDAO-2 may represent a dysfunctional OLs state that contributes to myelin loss and neuroinflammation in PD progression.

The negative correlation between MBP expression and the proportion of PDAO-1 and PDAO-2 underscores the critical role of OLs dysfunction in demyelination and neurodegeneration across neurological disorders, including PD^33,61,62,70–72^. The increase in these populations, along with the decline in MBP levels, suggests a transition in OLs functions away from myelination and towards alternative stress-related responses. This change may impair their ability to maintain and repair myelin sheaths, which further contributes to the degeneration of brain tissue and worsens neurodegenerative processes.

Additionally, two more subclasses are identified as PD-associated: PDAO-3 and PDAOPCs. PDAO-3, a mature oligodendrocyte subpopulation, exhibits transcriptomic alterations indicative of dual roles in stress adaptation and myelination. GSEA highlights upregulation in pathways associated with autophagy, sphingolipid metabolism, and cytokine signaling, suggesting PDAO-3 active engagement in cellular stress responses. However, these changes coincide with a marked downregulation of critical myelin assembly processes, including cell projection organization and integrin signaling. Spatial transcriptomics further demonstrates the enrichment of PDAO-3 in striatal white matter tracts, underscoring its potential role in the myelin deficits observed in PD^73,74^. This dual functionality of PDAO-3 may link stress responses to disrupted myelin maintenance and broader neuronal dysfunction in PD. PDAOPCs, a subset of OPCs, appear to play a significant role in early PD pathogenesis. These cells show heightened activity in pathways related to mitochondrial function and mitophagy, coupled with enhanced antigen presentation capabilities, indicating an adaptive response to cellular stress and metabolic demands in the PD microenvironment. However, the reduced expression of intercellular communication molecules, suggests a compromise in PDAOPCs ability to mature into myelinating oligodendrocytes. This impairment likely contributes to the observed decrease in myelin levels within the Pu of PD patients.

Additionally, PDAOPCs exhibit a distinct glutamate-related signaling profile, implicating them in altered neurotransmitter dynamics and interactions with neuronal circuits in PD. Together, these features point to PDAOPCs as an early, vulnerable oligodendroglia population whose dysfunction propagates broader myelin and connectivity deficits in PD.

Collectively, these findings highlight the need to unravel the mechanisms underlying OLs dysfunction in PD, as such insights are crucial for guiding the development of therapies that protect myelin and promote neuronal health. Moreover, the role of PDA populations in exacerbating neurodegeneration through the disruption of essential glial-neuronal interactions underscores their critical contribution to the dysfunction of the basal ganglia circuit that characterizes PD. Together, this growing body of knowledge not only emphasizes the interplay between glial and neuronal dysfunction in PD but also opens a promising avenue for the development of targeted therapeutic strategies aimed at addressing oligodendroglia pathology to slow or halt disease progression.

## Acknowledgements

The authors acknowledge the Massachusetts Alzheimer’s Disease Research Center (especially Patrick Dooley and Tessa Connors), the Parkinson’s UK Brain Bank at Imperial funded by Parkinson’s UK (a charity registered in England and Wales, 258197, and in Scotland, SC037554), and the NIH NeuroBioBank (NIH Ref-1851, Requestor: Professor Ernest Arenas, R.I.P.), for the supply of tissue samples and associated clinical and neuropathological data. The authors also want to thank the National Genomics Infrastructure in Stockholm funded by Science for Life Laboratory and SNIC/Uppsala Multidisciplinary Center for Advanced Computational Science for assistance with massively parallel sequencing and access to the UPPMAX computational infrastructure. Further thanks go to Katarina Tiklova and Chika Yokota from the *In Situ* Sequencing Infrastructure Unit at the Science for Life Laboratory, Stockholm. Finally, we are extremely grateful to the brain donors and their families who have made this study possible.

## Abbreviations

Ols: Oligodendrocytes
PD: Parkinson’s disease
OPCs: Oligodendrocyte Precursor Cells
COPs: Committed OPCs
MOLs: Mature Oligodendrocytes
CN: caudate nucleus
Pu: putamen
SNpc: Substantia Nigra pars compacta
LBs: Lewy bodes
LNs: Lewy neurites
AD: Alzheimer’s Disease
sc/sn-RNA-seq: single-cell/single-nucleus RNA-sequencing
MS: multiple sclerosis
GWAS: Genome Wide Association Studies
GSEA: Gene Set Enrichment Analysis
NES: Normalized Enrichment Score
FDR: False Discovery Rate
GO: Gene Ontology
KEGG: Kyoto Encyclopedia of Genes and Genomes
Pu: Putamen
CN: Caudate nucleus
PDA-OLs: PD-associated oligodendrocytes
GCN: graph convolutional network
WM: white matter
GM: gray matter
UMIs: unique molecular identifiers

