## supplementary figures for "Oligodendroglia vulnerability in the human dorsal striatum in Parkinson’s disease"

### Supplementary Material

#### Supplementary Figure 1

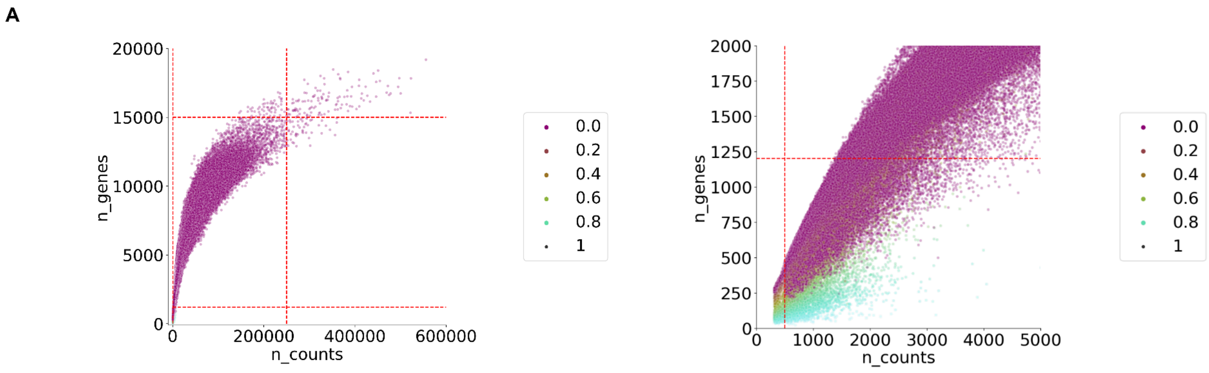


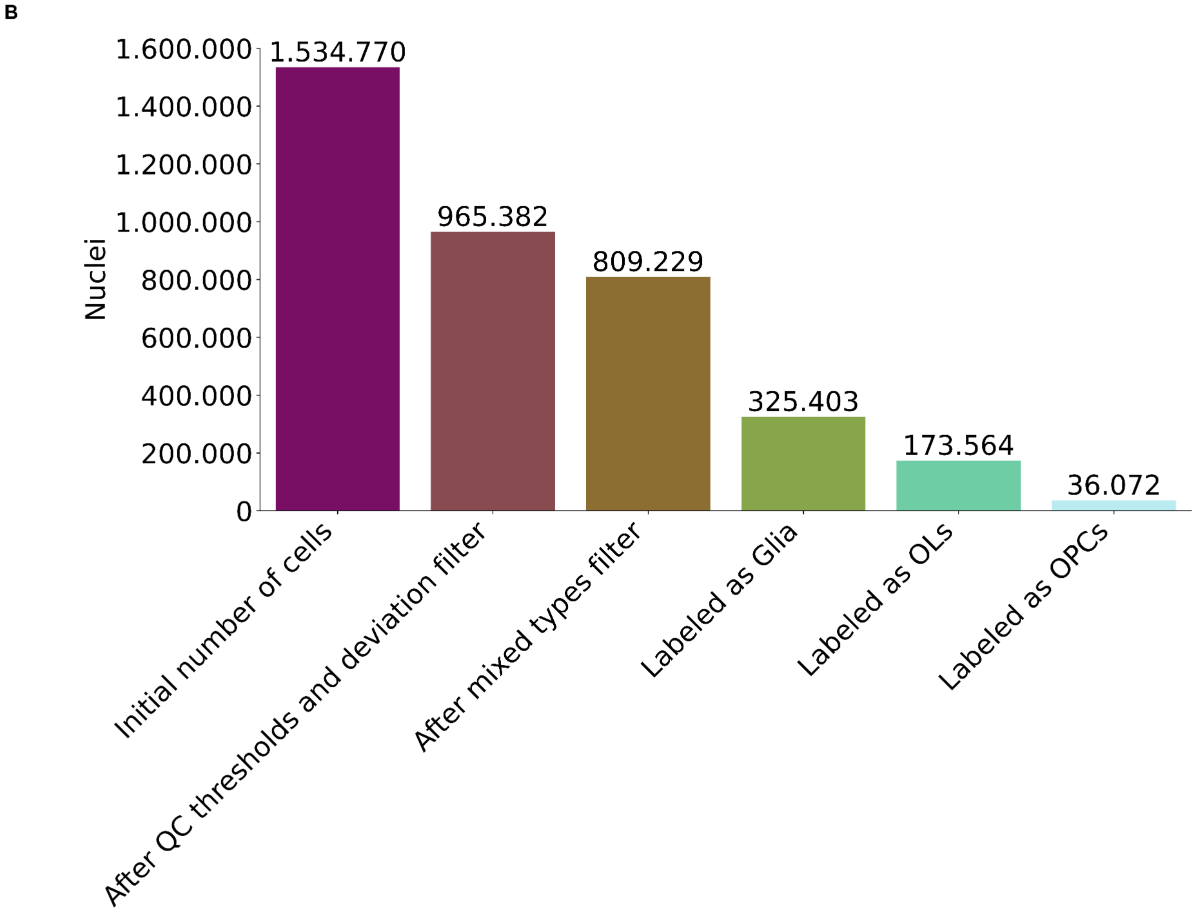

### Supplementary Figure 2.

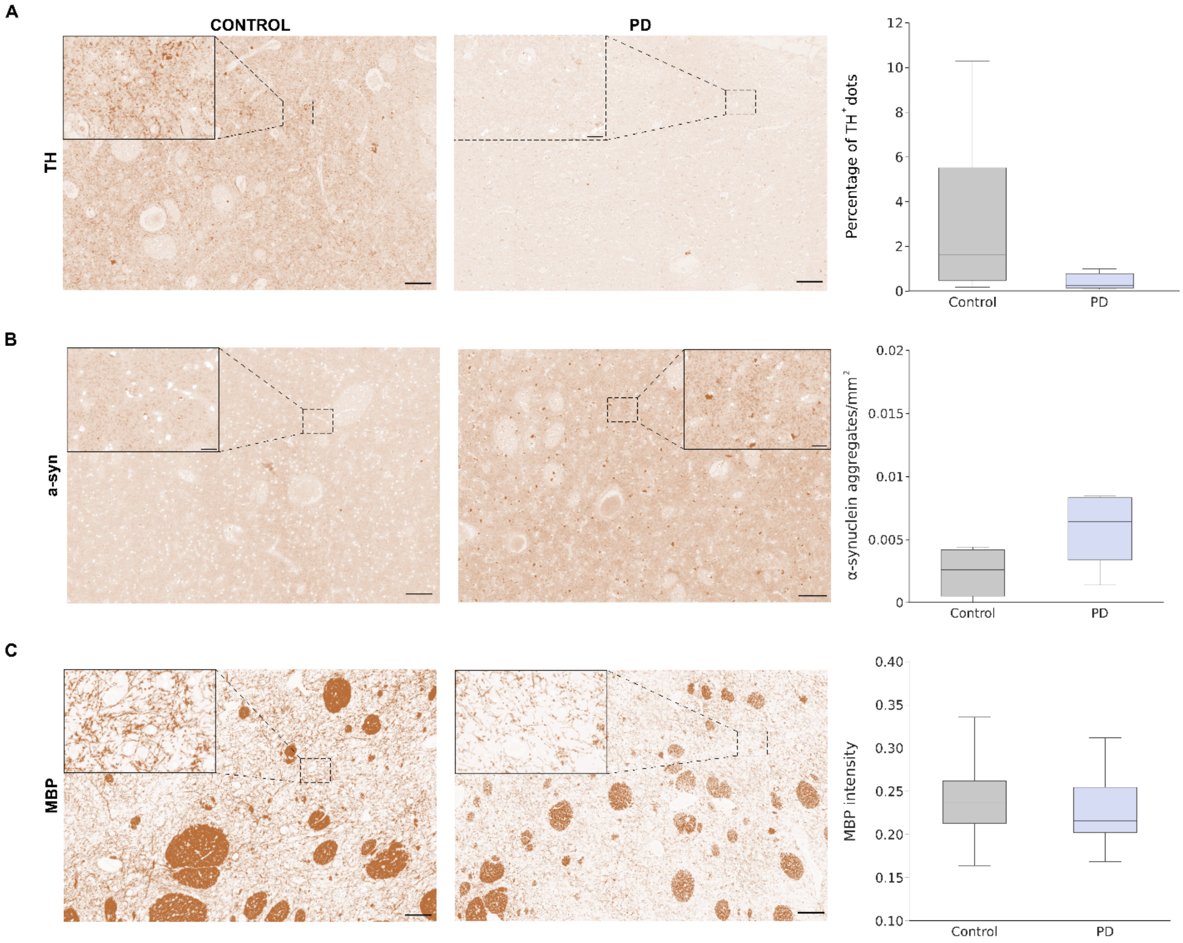

### Supplementary Figure 3

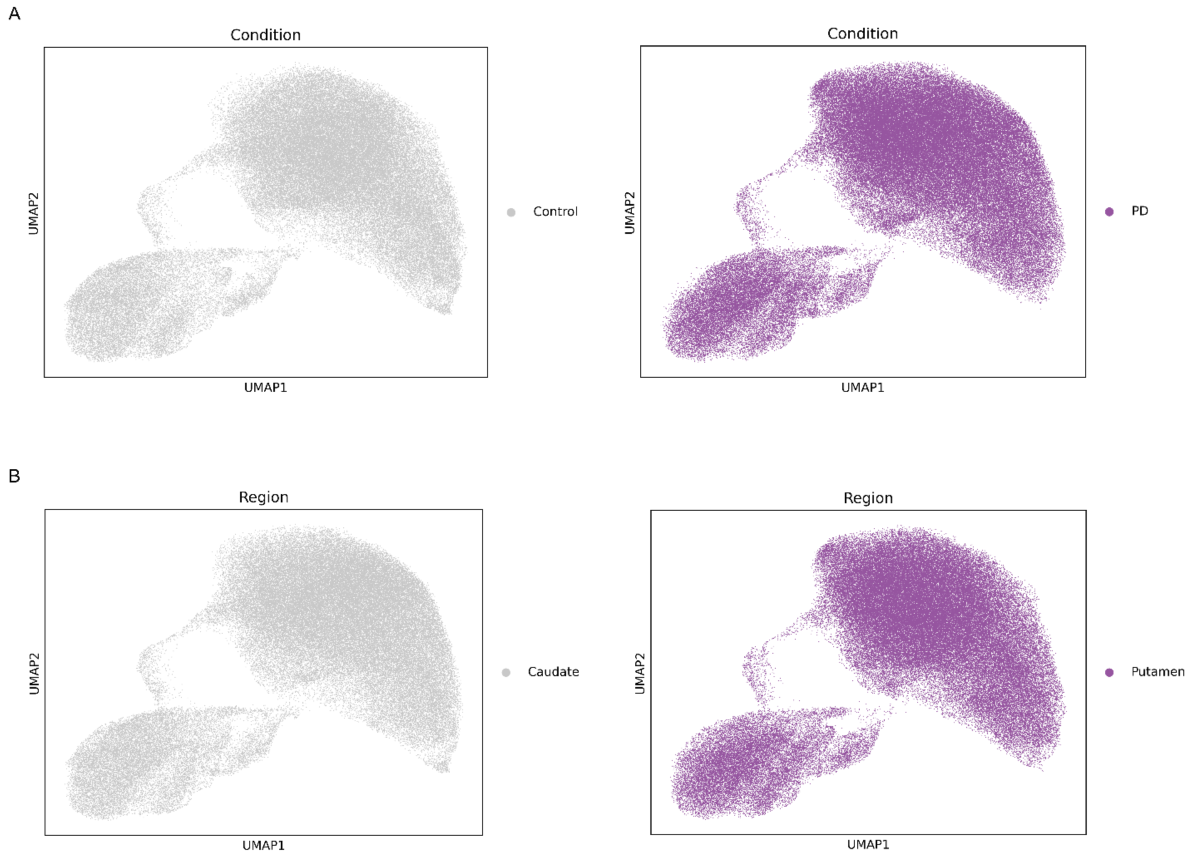

### Supplementary Figure 4

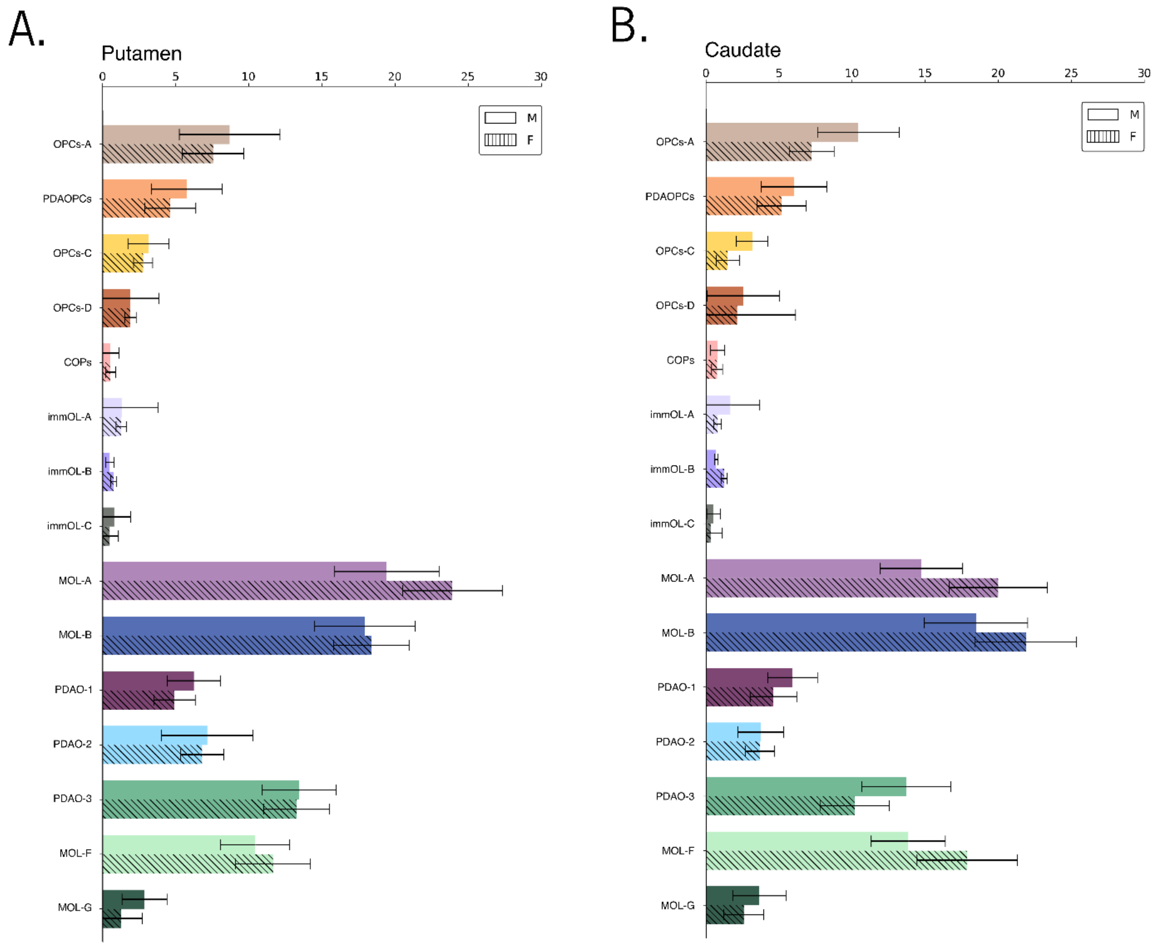

### Supplementary Figure 5

**A**

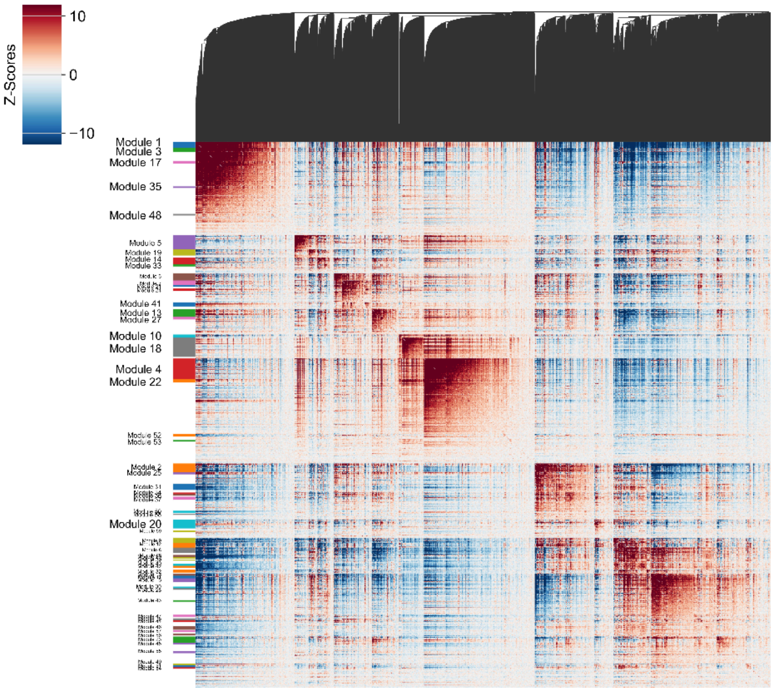


**B**

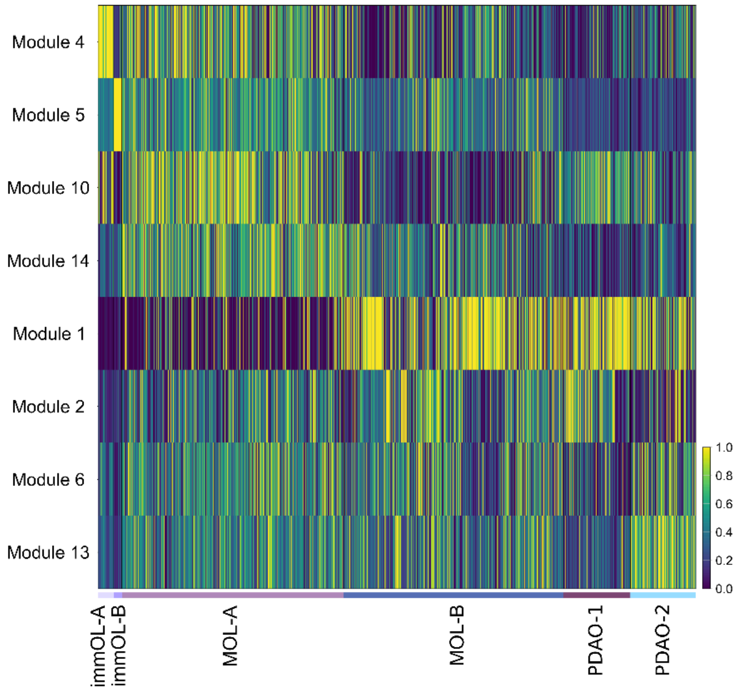

### Supplementary Figure 6

A

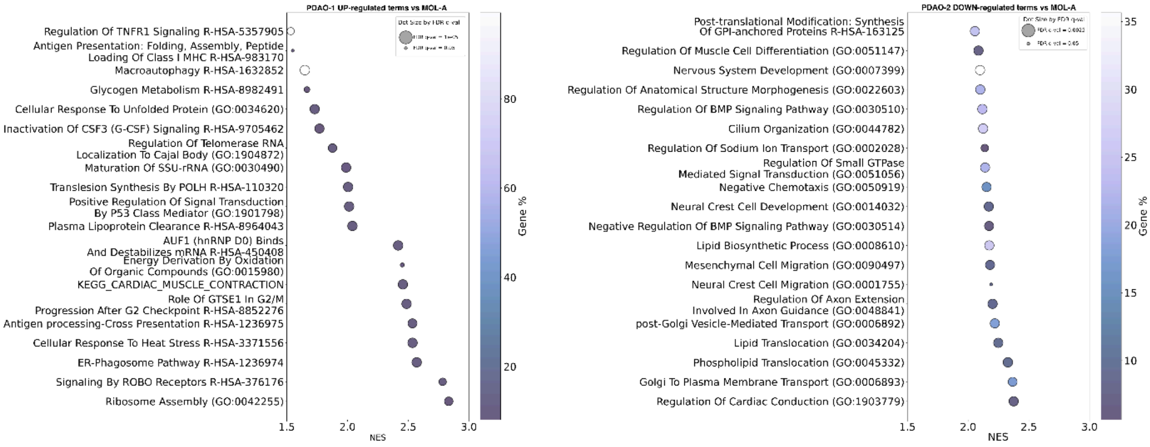


B

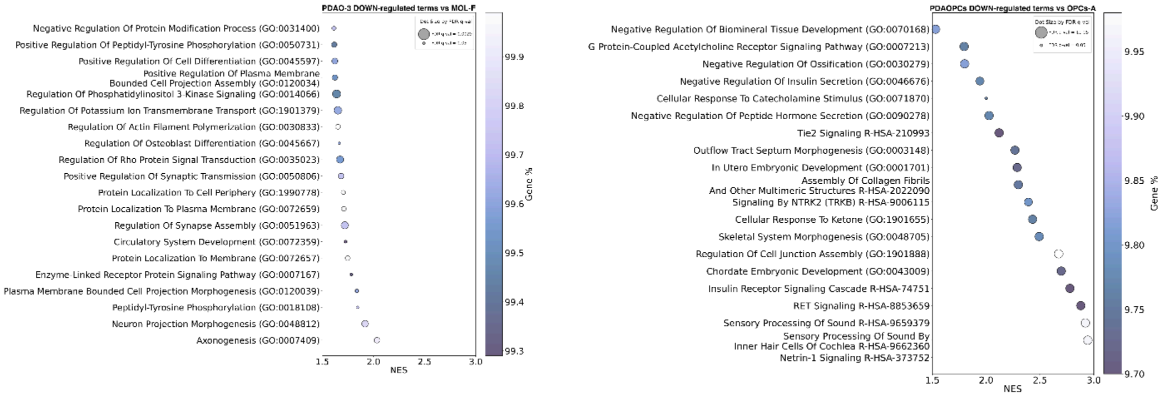

### Supplementary Figure 7

**A**

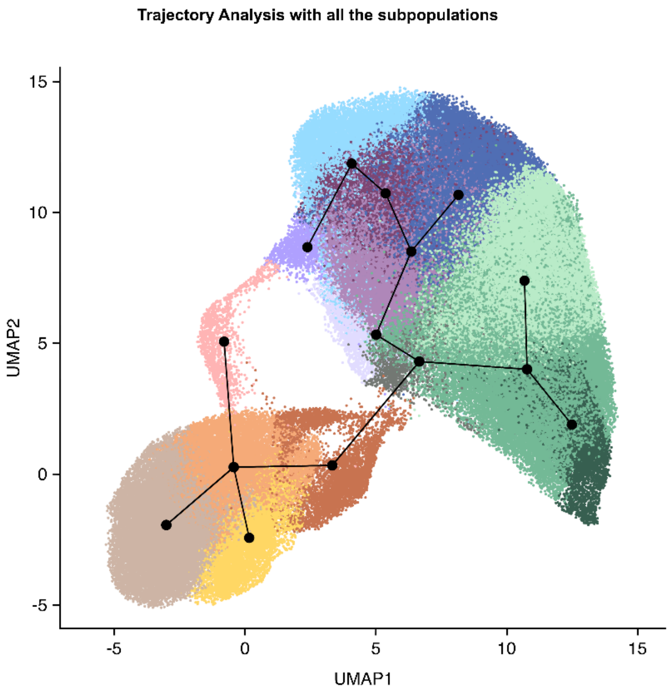


**B**

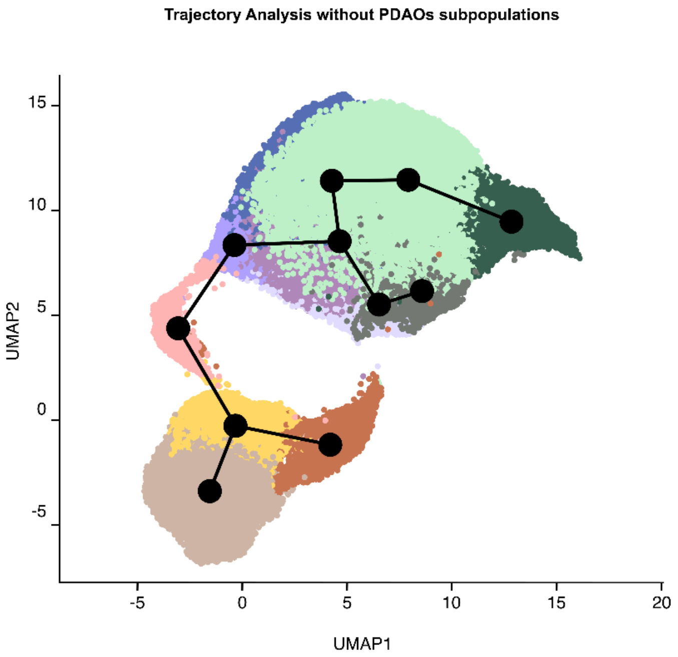

**Supplementary Figure 1. Quality control metrics for the snRNA-seq dataset. A.** Scatter plots with the distribution of gene counts and total counts per nucleus, stratified by the proportion of mitochondrial gene-derived UMIs among the total detected UMIs. Red lines indicate the values selected for the initial thresholding criteria (refer to Methods for details). **B.** Summary of the number of nuclei retained at each stage of the analytical workflow.

**Supplementary Figure 2. Immunohistochemical Analysis of Tyrosine Hydroxylase, α-Synuclein, and MBP in CN Region. A.** Immunohistochemical analysis of Tyrosine Hydroxylase in the CN region. Violin plots illustrate the distribution of marker expression in Control and PD conditions. **B.** Immunohistochemical analysis of α-synuclein in the CN region. Violin plots illustrate the distribution of marker expression in Control and PD conditions. **C.** Immunohistochemical analysis of MBP in the CN region. Violin plots illustrate the distribution of marker expression in Control and PD conditions. Statistical significance was determined using t-tests, with p-values indicated ($< 0.05$).

**Supplementary Figure 3. Cell Distribution Across Conditions and Regions. A.** UMAPs display the uniform distribution of cells from each condition across all subpopulations (Control – Left, PD – Right). **B.** UMAPs display the uniform distribution of cells from each region across all subpopulations (Caudate – Left, Putamen - Right).

**Supplementary Figure 4. Subpopulation Percentages by Sex and Region. A.** Barplots showing the percentages of each subpopulation, calculated based on the total number of nuclei depending on the sex and Pu area. **B.** Barplots displaying percentages of each subpopulation, calculated based on the total number of nuclei depending on the sex and CN region.

**Supplementary Figure 5. Hotspot Modules and Marker Genes Associated. A.** Heatmap illustrates modules detected by Hotspot analysis, based on local correlations within the dataset. **B.** Heatmap representing the association of modules with specific marker genes for each subpopulation, highlighting the distinct genetic signatures of the identified subpopulations.

**Supplementary Figure 6. GSEA Enriched Terms in PDAO Subtypes. A. (Left)** GSEA results highlighting the top 20 enriched terms (from GO, KEGG, and Reactome databases) with increased expression in PDAO-1 compared to MOL-A, ranked by gene ratio. (**Right**) Top 20 enriched terms with decreased expression in PDAO-2 relative to MOL-A, also ranked by gene ratio. **B. (Left)** Top 20 enriched terms showing down-regulation in PDAO-3 compared to MOL-F, based on gene ratio. (**Right**) Down-regulated terms (top 20) in PDAOPCs relative to OPCs-A, ranked by gene ratio.

**Supplementary Figure 7. Slingshot Pseudotime Analysis of Differentiation Trajectories Across Subpopulations. A.** Pseudotime UMAP generated via Slingshot including all the subpopulations. Each dot is related to a specific subpopulation, while lines indicate how the differentiation process occurs. **B.** UMAP visualization of differentiation trajectory excluding PDA clusters.
